# Hereditary Fusion Genes Are Associated with the Inheritance of Acute Myeloid Leukemia

**DOI:** 10.1101/2022.10.31.514615

**Authors:** Fei Ling, Degen Zhuo

**Author notes:** Correspondence should be addressed to: SplicinagCodes, BioTailor Inc, 7328 SW 82^nd^ Street, #B114, Miami, FL 33143 USA,.

## Abstract

Fusion genes are thought to be somatic and cause cancer, including acute myeloid leukemia (AML). Validating highly-recurrent fusion genes in healthy samples compelled us to systematically study hereditary fusion genes (HFGs). Here, we used curated HFGs to analyze AML fusion genes, and we identified 243 HFGs associated with AML inheritance from 926 potential HFGs. Many HFGs were one-to-many and many-to-one fusions that augmented signals from environmental and genomic alterations and seemed to support the “two-hit” hypothesis. The most highly-recurrent HFGs were also observed in multiple myeloma and monozygotic twin datasets, suggesting that AML is a complex genetic disease. HFGs, as cancer genetic biomarkers, are the most basic foundations for future genetic and genomic studies.

---

Acute myeloid leukemia (AML) is a type of blood cancer characterized by uncontrolled clonal expansion of hematopoietic progenitor cells. Cytogenetic profiling and molecular screening found that acquired recurrent genomic abnormalities aggravate malignant cell growth^1^. Advances in next-generation genome-wide sequencing and RNA-Seq have made it possible to identify gene mutations^2^ and genomic abnormalities^3^ resulting in fusion genes^4^. Even though familial clustering of AML cases has been known for more than five decades, several syndromes were shown to be syndromes with a genetic predisposition for AML development^5^. Although germline mutations associated with leukemia were discovered in more than 20 genes, they could explain only small percentages of these genetic predispositions^6^. The germline genetic lesions in a significant fraction of these families remain unknown^5-8^.

Fusion genes have been thought to be somatic and cancerous^9^. Recent works have shown that fusion genes were detected at high frequencies in noncancerous samples^10-17^. However, these apparent discrepancies were systematically removed from further consideration by researchers^18^ or software systems^19,20^ and hindered further investigations. We analyzed large numbers of RNA-Seq datasets and identified many fusion genes, among which *KANSARL* was validated as the first predisposition fusion gene specific to 29% of the European ancestry population^21^. Validating *TPM4-KLF2* and others in healthy samples compelled us to use a monozygotic (MZ) twin genetic model to systematically study hereditary fusion genes (HFGs), and we discovered 1180 HFGs^22^. HFGs were defined as the fusion genes that offspring inherited from their parents, excluding read-through transcripts^23^, which were named epigenetic fusion genes^22^. This report used curated HFGs to analyze fusion transcripts uncovered from 390 AML patients’ RNA-Seq data. We identified 243 HFGs, many of which formed one-to-many and many-to-one fusions and were genetic factors associated with AML inheritance.

We downloaded the RNA-Seq data of the Leucegene AML project^24^ from 390 AML patients. We used SCIF (SplicingCodes Identify Fusion Genes) to analyze the RNA-Seq data, and we identified 1,010,000 fusion transcripts, 114,000 of which were detected in 2 to 325 AML patients. Alternative splicing and repetitive sequences caused false positives, the rate of which was 1-2%^21^. Since the random genomic alteration rate was 3.6×10^-2 25^, the HFG frequency in a population would be 20-fold higher than the random genomic alteration rate^22^, distinguishing somatic abnormalities. We used 1180 HFGs previously discovered from MZ twins^22^ and searched for AML fusion transcripts. We identified 926 potential HFGsin1 to 325 AML patients (Fig.1a), which counted for 78.5% of 1180 MZ twins and much higher than 48.8% of the GTEx counterpart^22^, suggesting that these 926 HFGs had unfavorable AML risk profiles. To avoid confusion, we set a 10% recurrence frequency (RF) as the cutoff. Therefore, unless specified, HFGs with RFs of ≥10% were treated as HFGs or potential HFGs. We uncovered 243 HFGs with RFs ranging from 10% to 83.3%, with an average RF of 23.9% (Table S1). The number of HFGs identified was significantly larger than the number of germline mutations identified^7^, suggesting that HFGs were more critical genetic factors.

**Fig.1.**
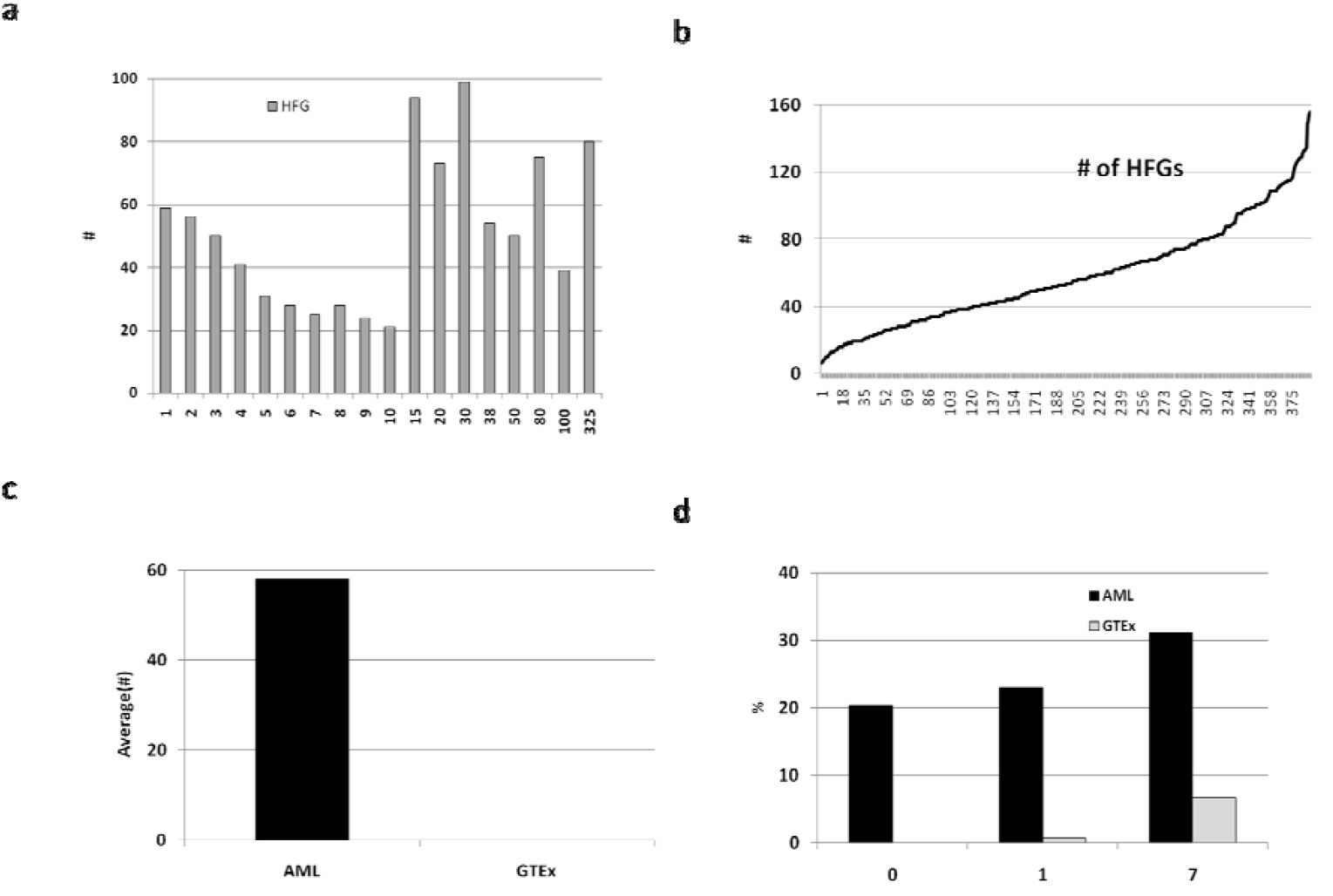
Identification of hereditary fusion genes (HFGs) associated with the inheritance of AML. a). Distribution of the recurrence frequencies of 926 potential HFGs. The dark gray rectangles represent the total number of potential HFGs. If increments were >1, the HFG numbers were summarized between two recurrence frequencies indicated on the horizontal axis; b). Distribution of the numbers of potential HFGs identified in 390 AML patients; c). Comparison of the average number of HFGs between AML and GTEx; d). Comparative analysis of recurrence frequencies of 243 HFGs in GTEx and AML. The solid black and gray rectangles represent AML and GTEx, respectively.

To better evaluate the results,427 healthy blood samples from GTEx were selected as controls. Similarly, we used the curated HFGs to analyze fusion transcripts from GTEx and uncovered 337 potential HFGs, which ranged from 0.23% to 41%, the average of which was 2.1%. Then, 66.8% of 243 AML HFGs were observed in the GTEx HFGs, supporting that HFGs were conserved and widespread^22^. Fig.1b shows that AML patients had from 6 to 156 HFGs, the average of which was 58- and 944-fold higher than the GTEx counterpart (Fig.1c), suggesting that these 243 HFGs were uncommon in healthy populations and were associated with AML. Fig.1d shows that as the RFs of GTEx HFGs increased, so did their AML counterparts, suggesting that most HFGs were under selection pressures. The GTExSQRDL*-B2M* RF was 27.4% and 26% higher than its AML counterpart (p≤0.005) and suggested that *SQRDL-B2M*wasthe only HFG conferring a favorable prognosis for AML. The statistical analysis showed that the RFs of the 240 AML HFGs were significantly higher than those of their GTEx counterparts, suggesting that these 240 HFGs were associated with AML (Table S1).

To characterize HFGs, we sorted 5’- and 3’-HFG gene IDs independently. Fig.2a shows that seven 5’-genes were fused with ≥5 3’-genes, thrice the 5’-gene average, suggesting that 5’-genes were not randomly distributed. *OAZ1* was the most recurrent 5’-fused gene and was fused with 24 different 3’-genes. Out of 11 *OAZ1*-*SPPL2B* fusion sites^23^, all main isoforms of these 24 5’-*OAZ1*-fused HFGs used the first *OAZ1* exon, which provided regulatory sequences for 3’-genes. Table S2 shows that eight, seven, five, and four 5’-*OAZ1*-fused HFGs encoded hybrid, truncated, intact, and short polypeptides, respectively, suggesting 5’-*OAZ1*-fused HFGs created diversities. Fig.2b shows that AML patients possessed up to 21 different *OAZ1*-fused HFGs, the average of which was 5.4. Multiple 5’-*OAZ1*-fused HFGs in an individual indicated that identical regulatory factors controlled these HFGs to form one-to-many fusions, potential natural networks. In contrast, Fig.2b shows that 52.5% of GTEx individuals had no single copy of 5’-*OAZ1*-fused HFGs, an average of which was 0.48 and 11-fold less than the former. Consequently, a single environmental factor or genetic alteration could affect all genes under this network and quickly initiate AML.

**Fig.2.**
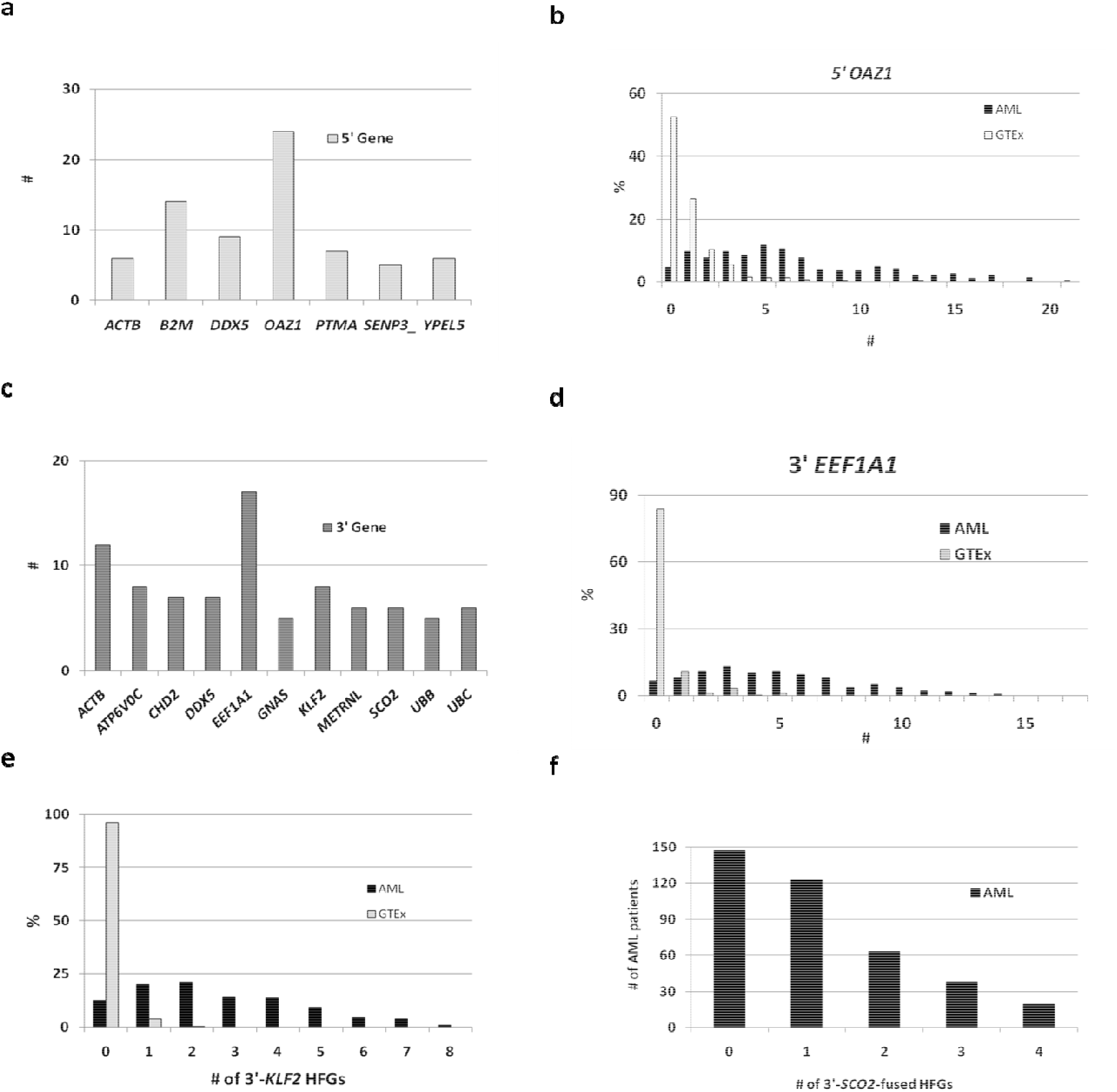
Characterization of 243 HFGs associated with AML inheritance. a). The list of 5’-genes fused with ≥5 3’-fused genes; b). The comparative analysis of 5’-*OAZ1*-fused HFGs among 390 AML patients and 427 GTEx individuals; c). The list of 3’-genes fused with ≥5 5’-fused genes; d). The comparative analysis of 5’-*EEF1A1*-fused HFGs among 390 AML patients and GTEx individuals; e). The comparative analysis of 3’-*KLF2*-fused HFGs between 390 AML patients and GTEx individuals; f).The distribution of 3’-*SCO2*-fused HFGs among AML patients. Solid black and gray rectangles represent AML and GTEx, respectively.

On the other hand, Fig.2c shows that eleven 3’-genes were fused with ≥5 5’-genes. The most recurrent 3’-gene was *EEF1A1*, fused with 17 different 5’-genes. The most recurrent isoforms of all 3’-*EEF1A1*-fused HFGs used the 5’-splice site of the second *EEF1A1* exon, located at the 5’-UTR of 3’-fused-*EEF1A1*.All 5’-genes provided their regulatory and 5’-UTR sequences, conferring only regulatory machinery. All 3’-*EEF1A1*-fused HFGs encoded intact eukaryotic translation elongation factor 1 α1, the overexpression of which was shown to inhibit p53- and p73-dependent apoptosis and chemotherapy sensitivity in cervical carcinoma cells^26^.Fig.2d shows that the numbers of 5’-genes fused with *EEF1A1* ranged from zero to 17, the average of which was 6.3. One individual’s maximum number of 5’-genes was 17, which formed many-to-one fusions that disrupted *EEF1A1* expression. In contrast, Fig.2d shows that 83.3% of GTEx individuals had no *EEF1A1*-fused counterparts, and the average was 0.013. The former was 484-fold higher than the latter and suggested that 3’-*EEF1A1*-involved HFGs were associated with AML.

Fig.2a&c shows that *ACTB* and *DDX5*, which encode β-actin and DEAD-box helicase 5, were fused with both 5’- and 3’-genes to form both one-to-many and many-to-one fusions. A 5’-*ACTB* and 3’-*ACTB* were fused with six different 3’-genes and twelve 5’-genes to form six 5’-*ACTB* HFGs and twelve 3’-*ACTB* HFGs. Three of six 5’-*ACTB* HFGs resulted in no protein alterations, while the remaining three altered N-termini; *ACTB-KLF2* had an ORF without a start codon (Fig.S1). Five 3’-*ACTB* HFGs encoded intact β-actin;*PTMA-ACTB,ELMO1-ACTB, KLF2-ACTB, TPM4-ACTB*, and *LFNG*-*ACTB* encoded β-actin-involved hybrid proteins; *TNRC18-ACTB* and *OAZ1-ACTB* coded for truncated β-actin (Fig.S2), which explained why *ACTB*, a housekeeping gene, was abnormally expressed in cancer^27^.*DDX5-CHD2, DDX5-EEF1A1, DDX5-HNRNPH3*, and *DDX5-UBB*encoded intact proteins, while *DDX5-ATP6V0C*, DDX5*-GNAS, DDX5-HNRNPU*, and *DDX5-SF1* produced truncated proteins (Fig.S3), which were from 43 to 702 aa shorter than their parental counterparts. Fig.2c shows that seven 5’-genes were fused with3’-*DDX5* and furnished their regulatory and 5’-UTRs.Since 3’-*DDX5* was fused in the coding region and had to use alternative start codon, *B2M-DDX5, HNRNPA2B1-DDX5, HNRNPH1-DDX5*, and *PTMA-DDX5* produced truncated proteins, while *OAZ1*-*DDX5* encoded a hybrid protein (Fig.S4). Hence, these 3’-*DDX5* HFGs changed *DDX5* gene regulation and encoded mutated DDX5 proteins. These one-to-many and many-to-one fusions might amplify environmental stresses and genomic abnormalities and quickly result in genetically heterogeneous clonal hematopoietic progenitors^28^. These HFGs disrupted the normal gene expression patterns, made cells more susceptible to environmental and genomic stresses, and provided a genetic basis for the two-hit hypothesis^29^.

To characterize HFGs, we used Morpheus to cluster these 243 HFGs.Fig.3 shows that HFGs were clustered into three groups: the highly-recurrent, moderately-recurrent, and sparsely-recurrent HFGs, with frequencies of 11.5%, 38.3%, and 50.2%, respectively. The highly-recurrent HFGs overlapped with the top 27 HFGs (Fig.1a), with RFs ranging from 40% to 83.3%.One of the most recurrent HFGs was *TPM4*-*KLF2*, which was detected in 74.1% of AML patients and encoded the tropomyosin 4-*kruppel*-like factor 2 hybrid protein. *TPM4*-*KLF2* was first reported in acute lymphoblastic leukemia (ALL)^30^.The validation of the high recurrence of *TPM4*-*KLF2*in multiple myeloma (MM) and healthy controls led us to study HFGs^22^. The MM *TPM4-KLF2* data were consistent with a previous report showing that *TPM4-KLF2* was found in 30% of AML samples and all three normal bone marrow samples^12^.

**Fig.3.**
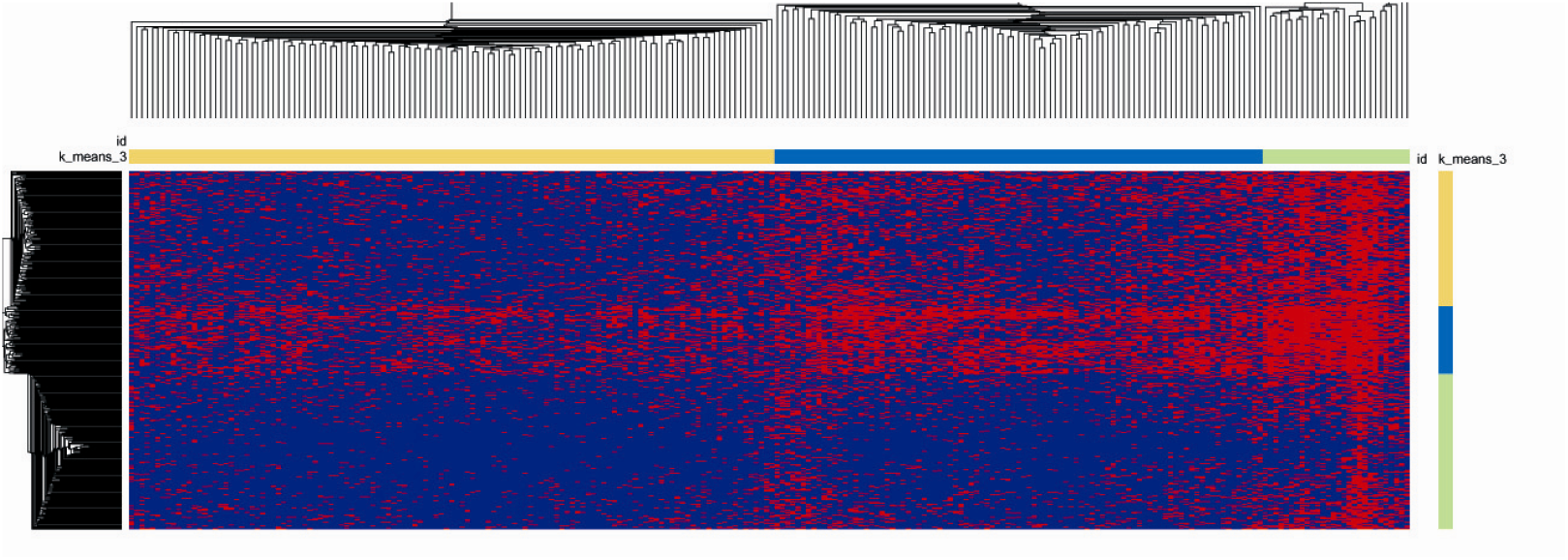
Heatmap of 243 HFGs among 390 AML patients generated by Morpheus. Horizontal light green, blue, and yellow rectangles represent the highly-recurrent, moderately-recurrent, and sparsely-recurrent HFGs. Vertical blue, yellow, and light green rectangles represent patients with highly abundant, moderately abundant, and sparsely-abundant HFGs, respectively.

Fig.2c shows that eight 5’-genes were fused with 3’-*KLF2* to form eight 3’-*KLF2*-fused HFGs and many-to-one fusions. Except for *OZA1*-*KLF2* without apparent ORFs, *ACTB-KLF2, AKAP8-KLF2, B2M-KLF2*, and *PTBP1-KLF2* resulted in frame shifts to produce ORFs similar to *kruppel*-like factor 2, while *TPM4-KLF2, ELL-KLF2*, and *EPS15L1-KLF2* were in-frame HFGs and produced hybrid proteins (Fig.S5). Fig.2e shows that 341 (87.4%) AML patients had from one to eight 3’-*KLF2*-fused HFGs, the average of which was 2.7, and 68.2% of 390 AML patients possessed two to eight 3’-*KLF2*-fused HFGs, suggesting potential network effects. In contrast, 96% of GTEx samples had no 3’-*KLF2*-fused HFGs (Fig.2e), suggesting that 3’-*KFL2*-fused HFGs were associated with AML.

Since *TPM4*-*KLF2* was observed in ALL^30^, AML^12^, healthy bone marrow^12^, MZ twin, and GTEx blood samples^22^, HFGs might not be disease-specific. We performed comparative analyses of HFGs from AML patients, MM patients, and MZ twins to study this notion. Table 1 shows that AML HFGs with RFs of ≥45% were also observed in MM patients and MZ twins. *TPM4*-*KLF2*, the most recurrent HFG, was detected in 74.4% of AML patients, 92.2% of MM patients, and 54.1% of MZ twins. The largest RF difference was found in*KRTAP3-1-POLK*, encoding truncated DNA polymerase kappa, which severely impaired the ability to allow the extension step of lesion bypass^31^; the *KRTAP3-1-POLK* RF was 12.9-fold higher in AML patients than in MM patients. Nevertheless, it was detected in 20.3% of MZ twins, which suggested that no single HFG could determine a phenotype. However, highly-recurrent HFGs supported the existence of diverse HFG genotypes, including some individuals who shared common HFGs for both MZ twin siblings to have ALL^32^, AML^33,34^, and Hodgkin’s disease^35^. These common HFGs provided genetic bases for diverse phenotypes from simultaneous AML and MM occurrences^36,37^, MM patients with an 11.51-fold increased risk of AML^38^ and second primary malignancies^39,40^.

**Table 1.** Recurrence frequency comparison of 15 HFGs among390 AML patients, 727 multiple myeloma (MM) patients, and 74 monozygotic (MZ) twins.

| HFGs | leukemia (399) | MM (727) | MZ (74) |
| --- | --- | --- | --- |
| *TRNAN35-FAM91A3P* | 81.5 | 56.4 | 32.4 |
| *C9orf100-DENND1C* | 72.9 | 17.7 | 44.6 |
| *TPM4-KLF2* | 72.4 | 92.2 | 54.1 |
| *OAZ1-KLF16* | 66.4 | 12.8 | 35.1 |
| *PTP4A1-EEF1A1* | 66.2 | 25.2 | 21.6 |
| *YWHAЕ-CRK* | 66.2 | 14.6 | 35.1 |
| *SYNCRIP-EEF1A1* | 61.9 | 12.2 | 17.6 |
| *BACH1-MECP2* | 55.4 | 15.8 | 58.1 |
| *CYTSB-CCDC144A* | 49.9 | 11.3 | 13.5 |
| *DDX5-EEF1A1* | 48.4 | 17.2 | 17.6 |
| *B2M-CHD2* | 48.4 | 14.0 | 43.2 |
| *YPEL5-FOSL2* | 48.4 | 10.0 | 8.1 |
| *KRTAP3-1-POLK* | 47.9 | 3.7 | 20.3 |
| *OAZ1-KLF2* | 46.9 | 58.2 | 33.8 |
| *NCOR2-UBC* | 46.6 | 32.3 | 35.1 |
| *HNRNPA2B1-DDX5* | 46.6 | 12.1 | 13.5 |

Fig.3 shows that based on individual HFG genotypes, AML patients were clustered into three groups: AML patients with highly-abundant, moderately-abundant, and sparsely-abundant HFGs (18.5%, 38.2%, and 43.3%, respectively). AML patients with highly-abundant HFGs had all types of HFGs, from sparsely-recurrent HFGs to highly-recurrent HFGs, suggesting that many HFGs functioned as groups. For example, *PIM3*-*SCO2, PLXNB*-*SCO2, PPP6R2*-*SCO2*, and *TRABD-SCO2*, were generated by local *SCO2* amplification^22^. Fig.2f shows that 30.8% of AML patients possessed ≥2 3’-*SCO2*-fused HFGs, and 19 AML patients had all four 3’-*SCO2*-fused HFGs. Therefore, these *SCO2*-fused HFGs were clustered into groups and were passed on to offspring. In contrast, Fig.3 shows that 43.3% of AML patients had sparsely-abundant HFGs. However, it was not known that these patients had sparsely-abundant HFGs for the following three reasons: the limited numbers of curated HFGs^22^, the technological limitation of our software system to discover fusion genes^21^, and 638 potential HFGs remained to be studied.

This work used the most fundamental principle of inheritance to study whether fusion genes were associated with AML inheritance. We used the curated HFGs to analyze fusion genes uncovered from AML patients, and we identified 243 HFGs associated with AML inheritance from 926 potential HFGs. Since the most recurrent AML HFGs were also found in MM patients and MZ twins, HFG genotypes could interact with the environmental stresses to promote random AML genomic abnormalities^21^ and produce diverse phenotypes. Hence, AML is a complex genetic disease and developmental interaction between environmental, germline, and somatic lesions. Future works must systematically validate and expand HFG databases and study interactions between HFGs and environmental factors. This work reveals only the tip of the HFG iceberg and presents the first line of evidence that cancers such as AML or MM are complex genetic diseases. HFGs, as the most fundamental genetic concept, marked the dawn of the digitalization of applied and theoretical genetics and genomics.

## Methods

### Materials

#### Human acute myeloid leukemia (AML) RNA-Seq dataset

RNA-Seq data of the Leucegene AML project^24^ (GEO: GSE67040) were downloaded from NCBI. We identified heparinized blood samples from 391 unique AML patients. We removed one sample with poor-quality RNA-Seq data from this study. Hence, this dataset included data from 390 unique AML patients.

#### Genotype-Tissue Expression (GTEx) blood RNA-Seq data

We selected GTEx blood samples as controls to better evaluate the results. We downloaded the GTEx blood RNA-Seq data (dbGap-accession: phs000424.v7.p2), from which we identified 427 healthy blood samples.

#### Human multiple myeloma RNA-seq dataset

MMRF CoMMpass RNA-seq data (phs000748.v5.p4) were downloaded from the controlled-access dbGap (https://www.ncbi.nlm.nih.gov/gap)^41^. We identified 727 newly diagnosed multiple myeloma patients with unique IDs. This work has been described previously (Ling and Zhuo, submitted).

#### Computers

All computations were performed on Linux Desktop computers with 8 to 12G memory.

#### Software

All analytic software was developed in-house and written in Perl programming.

## Methods

### Identification of fusion transcripts by SCIF (<u>S</u>plicing<u>C</u>odes <u>I</u>dentify <u>F</u>usion Transcripts

SCIF (<u>S</u>plicing<u>C</u>odes Identify <u>F</u>usion Transcripts) was developed in the C and Perl programming languages. The algorithm used by SCIF has been described in detail previously^21^. The most important features of this algorithm were that a splicingcode table was precomputation and that the minimum unit analyzed was a raw RNA-Seq read without any further assembly. A raw RNA-Seq read could not be further divisible and functioned like zero and one in computer science. The most critical parameter was the minimum sequence length of the 5’- and 3’-genes. We used the minimum length of a default length of 20 bp. At a sequence length of 20 bp, we could remove highly repetitive DNA sequences such as *ALU* and *LINE1* elements to increase the computation speed. At the same time, we could retain some moderately repetitive sequences and easily verify the analyzed results during the following validation experiments. The algorithm used the pre-computated splicing code table, and all products were products spliced from transcribed sequences with spliceosomal introns. Hence, we could predict that all false-positive artifacts were from repetitive sequences, pseudogenes, and alternatively-spliced products. We easily identified false positives by BLAST and other manual and computation methods. From the algorithm, SCIF could not discover the fusion products generated by tandem duplications and fusion products from the sequences which were not in the table.

### Identifying hereditary fusion genes (HFGs) in acute myeloid leukemia (AML)

To discover HFGs in AML, we first used SCIF to analyze the RNA-Seq data of the Leucegene AML project (GEO: GSE67040) at the default parameters and discovered 1.1 million fusion transcripts. Then, we determined the quality of the uncovered fusion transcripts by manually inspecting some fusion transcripts from *KANSARL* and epigenetic (read-through) fusion genes (EFGs). Then, we compared these raw fusion transcripts with those from 37 pairs of monozygotic (MZ) twins to ensure that the data were accurate and relatively comparable. After inspection, we used 1180 HFGs discovered and curated from a previous study of MZ twins ^22^to analyze the total fusion transcripts from 390 AML patients. We used 5’ 20 bp and 3’ 20 bp fusion junction sequences to discover fusion transcripts with identical 5’ and 3’ fusion junction sequences. If an RNA-Seq read had identical fusion junctions, this fusion transcript was thought to be from HFGs. Furthermore, we removed 67 fusion transcripts that overlapped with the fusion transcripts of MZ twin HFGs, with too many alternative fusion gene IDs and only one isoform. We identified a total of 926 potential HFGs. Since we could not distinguish the HFGs with low recurrence frequencies from somatic fusion genes, we set the recurrence frequency cutoff at 10%. Only potential HFGs with recurrence frequencies of ≥10% were treated as HFGs in this study.

### Recurrence frequencies (RFs)

The recurrence frequency (RF) was defined as the number of HFG-positive individuals divided by the total number of samples used in a study. To rule out the potential effects of different experiments and protocols, if a sample was found to have one fusion junction of an HFG, this sample was treated as HFG-positive regardless of the copy numbers of this HFG.

### Protein sequence alignments

Protein sequence alignments were performed by COBALT (Constraint-based Multiple Alignment Tool) at NCBI (https://www.ncbi.nlm.nih.gov/tools/cobalt/cobalt.cgi?LINK_LOC=BlastHomeLink).

The alignment results were saved in Nexus format.

### Generation of heatmap by Morpheus

We first produced the files of 390 AML patients possessing 243 HFGs with recurrence frequencies of ≥10% to generate a heatmap. Then, we used Morpheus (https://software.broadinstitute.org/morpheus/) from the Broad Institute. We first used k-means clustering with Euclidean distance and three members to cluster 243 HFGs and 390 AML patients. Then, we further clustered the HFGs and patients by hierarchical clustering. The heatmap was saved as a PDF file.

### Classifications of types of fusion transcripts

The types of fusion transcripts that were clustered have been described previously^22^. They included deletions, inversions, intrachromosomal and interchromosomal translocations, and read-throughs.

### Distinguishing fusion transcripts caused by genomic alterations from *trans-*spliced fusion transcripts

It had been known that fusion transcripts might be *trans-*spliced products. Based on the splicingcode theory, *trans-*splicing and *cis*-splicing have an identical mechanism to remove spliceosomal introns but at much lower efficiencies. *Trans*-splicing may generate many fusion transcripts. However, our observation suggested that the gene expression levels of *trans*-spliced transcripts may be much lower than those of *cis*-spliced transcripts. We estimated that most fusion transcripts were from genomic alterations. In particular, it was mathematically impossible for fusion transcripts generated by *trans*-splicing to have more than one copy. Therefore, it was not plausible that the data presented in this study were due to *trans*-splicing.

### Statistical analysis

To know whether there were differences between the two datasets, we used Z-test to perform statistical analysis. Z-scores were calculated based on the following formula:

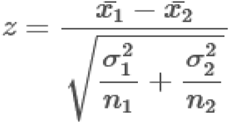

When we performed comparative HFG recurrent frequencies between AML and GTEx datasets, the HFGs were not present in the GTEx counterparts. If they were in these situations, we assumed that the GTEx counterparts had the minimum values observed in the AML dataset. *KANSARL* (*KANSL1-ARL17A*) was previously present in 28.9% of healthy European ancestral populations and associated with many types of cancer. However, the *KANSARL* recurrent frequency in GTEx was 38.4%, much higher than 28.9% of healthy European ancestral populations. Precautions must be taken care of.

## Acknowledgments

We express our most profound appreciation to Ms. Xiaoyan Yang, Prof. Benoit Chabot, Prof. Jeff Xiwu Zhou, Prof. Shunbin Ning, Prof. Yanbin Zhang, Prof. Yinxiong Li, and Mr. Noah Zhuo for their various contributions and support during the last two decades.

