## Supplementary Table 1 for "Hereditary Fusion Genes Are Associated with the Inheritance of Acute Myeloid Leukemia"

### Supporting Materials

**Contents**

**Table S1** ……………………………………………………………………………………………………………..2 -- 6

**Table S2** ……………………………………………………………………………………………………………...7 -- 7

**Fig.S1** ……………………………………………………………………………………………………………….9 -- 11

**Fig.S2** ………………………………………………………………………………………………………………..12 -- 14

**Fig.S3** ………………………………………………………………………………………………………………..15 -- 20

**Fig.S4** ………………………………………………………………………………………………………………...21 -- 23

**Fig.S5** …………………………………………………………………………………………………………………24 -- 24

### Table S1 Comparative analysis of 243 HFGs between AML and GTEx datasets

| HFG IDs | 5' Junction Sequences | 5' Junction Sequences | AML (%) | GTEx (%) | Folds | *p* |
|---|---|---|---|---|---|---|
| *TPM4-KLF2* | AGCGCGAGCGGCGCGAGAAA | GTGAGAAGCCCTACCACTGC | 74.1 | 0.9 | 269.4 | **** |
| *TRNAN35-FAM91A3P* | TGAGGACTGCCTGTTTGCAG | GCTCTGATTGTACTTTCTCC | 83.3 | 13.1 | 210.3 | **** |
| *PTP4A1-EEF1A1* | CAATGAGTAAACATATTCCT | GTGTCGTGAAAACTACCCCT | 67.7 | 1.4 | 210.3 | **** |
| *SYNCRIP-EEF1A1* | CGCCTCAGCCCAACGGGGAG | GTGTCGTGAAAACTACCCCT | 63.3 | 0.2 | 208.1 | **** |
| *YWHAE-CRK* | GCAGGCTGAGCGATACGACG | GGGTGAGCCCCTCCAGACTC | 67.7 | 2.8 | 202.6 | **** |
| *OAZ1-KLF16* | GGCGGCAGCAGCAGTGAGAG | GGGAACGCCCTTTTGCTTGT | 67.9 | 5.2 | 195.0 | **** |
| *DDX5-EEF1A1* | GACCGCGGCCGGGACCGAGG | GTGTCGTGAAAACTACCCCT | 49.5 | 0.2 | 189.5 | **** |
| *B2M-CHD2* | TGGCCTGGAGGCTATCCAGC | GTGACTTAAGAGATTAAAAT | 49.5 | 0.2 | 181.8 | **** |
| *KRTAP3-1-POLK* | CCCATTGATTCTACATTATG | TCTACTTCAAATTACCATGC | 49.0 | 0.2 | 175.3 | **** |
| *CYTSB-CCDC144A* | TAGGCATCAAACCCAGCCTG | CATTCAGCAACTAGTTCCTG | 51.0 | 1.4 | 155.6 | **** |
| *YPEL5-FOSL2* | GGATCTCACCGCCGCTCAGG | AAATTCCGGGTAGATATGCC | 49.5 | 0.9 | 150.1 | **** |
| *HNRNPA2B1-DDX5* | CGACTGAGTCCGCGATGGAG | GTTTGGTGCACCTCGATTTG | 47.7 | 0.2 | 149.0 | **** |
| *BACH1-MECP2* | AATCAGAAATTGAGAAGCTG | ACGGAGTCTTGCTCTGTCAC | 56.7 | 4.9 | 149.0 | **** |
| *KLF10-AZIN1* | CCTCTCTCCAGCAGACTGCG | GGCGAACTTTCTGACCAAGT | 44.6 | 0.2 | 147.9 | **** |
| *TEX2-DDX5* | AGGCAGCCAGCGGATCTTGG | GTTTGGTGCACCTCGATTTG | 45.9 | 0.2 | 147.9 | **** |
| *OAZ1-KLF2* | GGCGGCAGCAGCAGTGAGAG | GTGAGAAGCCCTACCACTGC | 47.9 | 2.1 | 144.6 | **** |
| *BCLAF1-EEF1A1* | ACCGCGATTCGGCGTGTCAG | GTGTCGTGAAAACTACCCCT | 42.8 | 0.2 | 143.5 | **** |
| *UBQLN1-HNRNPK* | AGAATAGCTCCGTCCAGCAG | GCTACTGCAGCACTGGGGTG | 43.6 | 0.9 | 134.8 | **** |
| *KLF10-YWHAZ* | CCTCTCTCCAGCAGACTGCG | AACATCCAGTCATGGATAAA | 41.3 | 0.2 | 133.7 | **** |
| *NHP2L1-LLPH* | TCGGAAAGGAGCCAATGAGG | GTAAAACATGGCTAAAAGCT | 43.1 | 2.6 | 127.1 | **** |
| *SPG21-NEK9* | AGGTCAATCTTTATGTACAG | AGTTTCGCTCTTGTTGCCCA | 45.9 | 3.0 | 127.1 | **** |
| *RHOA-LOC100499183* | ACTCGGATTCGTTGCCTGAG | AGATGGTGTCTTGCTATGTT | 42.1 | 2.1 | 109.6 | **** |
| *ACTB-KLF2* | CCGTCCACACCCGCCGCCAG | GTGAGAAGCCCTACCACTGC | 37.7 | 0.5 | 107.4 | **** |
| *OAZ1-BTBD2* | GGCGGCAGCAGCAGTGAGAG | GTTCGTGCTGGCCGTGGGCA | 39.2 | 1.2 | 103.0 | **** |
| *OAZ1-DOT1L* | GGCGGCAGCAGCAGTGAGAG | GTAACTAGGATTTCTACCTC | 36.9 | 0.5 | 101.9 | **** |
| *HNRNPA2B1-EEF1A1* | CGACTGAGTCCGCGATGGAG | GTGTCGTGAAAACTACCCCT | 36.7 | 0.2 | 101.9 | **** |
| *AKAP8-KLF2* | CATGGACCAGGGCTACGGAG | GTGAGAAGCCCTACCACTGC | 35.1 | 0.2 | 98.6 | **** |
| *YWHAZ-AZIN1* | ACCACCCACTCCGGACACAG | GGCGAACTTTCTGACCAAGT | 35.4 | 0.2 | 98.6 | **** |
| *KIAA0182-KLHL36* | GCCTCTGTATTACTTACCAG | GGCTGAAATCTCTTTAATGA | 34.9 | 0.2 | 97.5 | **** |
| *ACTB-MAFK* | CCGTCCACACCCGCCGCCAG | CTACGAGTTCCAGGGAGCTC | 35.1 | 0.2 | 97.5 | **** |
| *PTMA-ACTB* | GCTCCGAAATCACCACCAAG | CTCACCATGGATGATGATAT | 34.1 | 0.2 | 96.4 | **** |
| *HNRNPH1-DDX5* | CTTGCAGTTCAGCGACCACG | GTTTGGTGCACCTCGATTTG | 34.9 | 0.2 | 96.4 | **** |
| *PPP6R2-SCO2* | GAAGCCAACGACCACACGCA | GAGCATCAGATCCATGCTGC | 37.9 | 2.8 | 95.3 | **** |
| *SYNCRIP-PNRC1* | CGCCTCAGCCCAACGGGGAG | GTTTTAAAATCAAAGATGGG | 33.8 | 0.2 | 95.3 | **** |
| *TTYH3-MAFK* | AGGACACCGACTACCAGCAG | CTACGAGTTCCAGGGAGCTC | 31.5 | 0.2 | 94.3 | **** |
| *UBR5-AZIN1* | GAGGACCAGCTCAATGACAG | GGCGAACTTTCTGACCAAGT | 31.8 | 0.2 | 94.3 | **** |
| *TRAF6-DYM* | TGGCATTTGAAGATAAAGAG | AGTTTCACTCTTGTTGCCCA | 38.7 | 3.7 | 93.2 | **** |
| *SEPT2-ERI2* | GTTCACACTGATGGTGGTCG | AAAGGGTACACTCGCCAGCA | 34.1 | 1.6 | 91.0 | **** |
| *DDX5-UBB* | GACCGCGGCCGGGACCGAGG | GTCAAAATGCAGATCTTCGT | 33.1 | 1.2 | 91.0 | **** |
| *OAZ1-LRCH4* | GGCGGCAGCAGCAGTGAGAG | CCCACAGCCCCCATCCCGTA | 31.5 | 0.5 | 91.0 | **** |
| *YARS2-NAP1L1* | AGAATGGACTAGGACTCAGT | GGTCTACCTTCTGCTTCCCT | 34.9 | 2.3 | 89.9 | **** |
| *FAM60A-UBC* | GGAACGGCTGCTTTTGGAAG | ACAATGCAGATCTTCGTGAA | 32.3 | 1.2 | 88.8 | **** |
| *ARHGAP26-NR3C1* | ACTCATAAGCGCGCTCAAGA | TTGATATTCACTGATGGACT | 30.0 | 0.2 | 87.7 | **** |
| *LRRC37A4-LRRC37A3* | TACCTACAGCAGCACTCCAG | GTCAGTTGCTCTCCCTGGAA | 30.0 | 0.2 | 86.6 | **** |
| *PHF3-PTP4A1* | CGCTCGTCTGGCGGAGCTGG | CAATTCTGTGGTGTTCTTGG | 31.5 | 0.9 | 85.5 | **** |
| *UBE2L3-SPOPL* | AGACCATCCTGGCTAACACG | GTGAGTGTGGTCCAAGATTC | 30.3 | 0.5 | 84.4 | **** |
| *OAZ1-ATP6V0C* | GGCGGCAGCAGCAGTGAGAG | CCCTGGGCGCTGCCTATGGC | 33.6 | 1.9 | 84.4 | **** |
| *B2M-ATP6V0C* | TGGCCTGGAGGCTATCCAGC | CCCTGGGCGCTGCCTATGGC | 30.3 | 1.2 | 83.3 | **** |
| *AMD1-EEF1A1* | GATCTTCGCACTATCCCAAG | GTGTCGTGAAAACTACCCCT | 29.2 | 0.5 | 83.3 | **** |

| | | | | | | |
|---|---|---|---|---|---|---|
| *ST3GAL1-YWHAZ* | CCAGCGTGTCTCCAGCCGTG | AACATCCAGTCATGGATAAA | 29.2 | 0.5 | 83.3 | **** |
| *GUK1-ARF1* | GGGCCGGGCCCCACCGGACG | GTGTCCCTGGCCAGTGTCCT | 45.4 | 10.1 | 82.2 | **** |
| *PLXNB2-SCO2* | CCTTGGCGGCGGACGTGCAG | GAGCATCAGATCCATGCTGC | 32.6 | 2.3 | 82.2 | **** |
| *RNPS1-ATP6V0C* | GGTGGCGGCGGCGGGAAGAT | CCCTGGGCGCTGCCTATGGC | 25.9 | 0.2 | 81.1 | **** |
| *FHL3-IFNAR2* | TGGGGAGACTGTCATGCCTG | TAGAGACGGGGTTTCACCAT | 31.0 | 2.3 | 81.1 | **** |
| *OAZ1-ZBTB7A* | GGCGGCAGCAGCAGTGAGAG | GTCTCGGCGCGGAAGATGGC | 29.2 | 1.4 | 81.1 | **** |
| *SLC19A1-SUMO3* | GCCTCCCGGTCCGCAAGCAG | GAGGGTGTGAAGACAGAGAA | 26.4 | 0.5 | 79.5 | **** |
| *PRPS1-GATAD2B* | CTGAGTAGCTGGGATTACAG | GATGGATAGAATGACAGAAG | 26.2 | 0.5 | 78.9 | **** |
| *NCRNA00115-FBXO25* | CGGAGCCAAGGTCCGCTCGG | AGGCATGGCTATTGCACCTT | 37.9 | 6.3 | 78.9 | **** |
| *AZIN1-YWHAZ* | TCTTTGGGCCGTTATCTCAC | AACATCCAGTCATGGATAAA | 26.4 | 0.5 | 78.9 | **** |
| *PTMA-ING5* | GCTCCGAAATCACCACCAAG | GTATCGAGAACCTTCCCTGC | 25.4 | 0.2 | 78.1 | **** |
| *TRNAN35-SRGAP2P2* | TGAGGACTGCCTGTTTGCAG | GCAAGTGAACCGGAGCAAAC | 27.2 | 1.4 | 77.8 | **** |
| *TBC1D25-USP4* | CCAAGTAGCTGGGATTACAG | ATCCTGAGAGTCAGACCTTG | 27.4 | 1.2 | 77.8 | **** |
| *OAZ1-DAZAP1* | GGCGGCAGCAGCAGTGAGAG | GAAGCTCTTCGTGGGCGGTC | 26.4 | 1.2 | 76.7 | **** |
| *OAZ1-CCNG2* | GGCGGCAGCAGCAGTGAGAG | ATGAAGGATTTGGGGGCAGA | 24.4 | 0.2 | 76.7 | **** |
| *PTMA-DDX5* | GCTCCGAAATCACCACCAAG | GTTTGGTGCACCTCGATTTG | 24.1 | 0.2 | 75.6 | **** |
| *MRPL20-SSU72* | TCATTGGGAATTTAGTTAAG | CAAACGGGGATTCAGCGTCC | 24.1 | 0.2 | 73.5 | **** |
| *OAZ1-LIMD2* | GGCGGCAGCAGCAGTGAGAG | AACCCAGCGGGTGCCGCTTC | 25.6 | 1.2 | 71.3 | **** |
| *PRPS1-BACH1* | CTGAGTAGCTGGGATTACAG | CAAAGTGAAAAGGAGAGCTT | 27.4 | 1.9 | 71.3 | **** |
| *HNRNPH1-EEF1A1* | CTTGCAGTTCAGCGACCACG | GTGTCGTGAAAACTACCCCT | 23.3 | 0.2 | 70.2 | **** |
| *ACTB-EEF1A1* | CCGTCCACACCCGCCGCCAG | GTGTCGTGAAAACTACCCCT | 23.3 | 0.2 | 69.1 | **** |
| *LOC729126-RABGEF1* | TTCCACATCACTGCAATAAA | AAAACTGTGGATTTAGTTAC | 32.6 | 4.7 | 68.0 | **** |
| *OAZ1-DDX5* | GGCGGCAGCAGCAGTGAGAG | GTTTGGTGCACCTCGATTTG | 23.1 | 0.2 | 68.0 | **** |
| *B2M-DDX5* | TGGCCTGGAGGCTATCCAGC | GTTTGGTGCACCTCGATTTG | 23.1 | 0.2 | 68.0 | **** |
| *PTMA-EEF1A1* | GCTCCGAAATCACCACCAAG | GTGTCGTGAAAACTACCCCT | 24.9 | 0.9 | 68.0 | **** |
| *RAB35-UBC* | GCTCATCATCGGCGACAGCG | ACAATGCAGATCTTCGTGAA | 24.1 | 0.7 | 66.9 | **** |
| *SENP3_-EEF1A1* | TCTGCGAGCCAGGATTCCCG | GTGTCGTGAAAACTACCCCT | 22.8 | 0.2 | 66.3 | **** |
| *CPSF6-DYRK2* | TCGGCGAAGAGTTCAACCAG | ATTGGCGGCAGTAAGCACAC | 22.8 | 0.2 | 65.8 | **** |
| *POLR2J4-ALKBH4* | CTGGGAAACATCATTAAATC | AAAACATACCGTTTCATTTA | 22.6 | 0.2 | 64.7 | **** |
| *OAZ1-CSNK1G2* | GGCGGCAGCAGCAGTGAGAG | CTGTGAGCCGTGAGCTTTGA | 30.0 | 3.5 | 64.7 | **** |
| *OAZ1-PTBP1* | GGCGGCAGCAGCAGTGAGAG | CGGGGATCTGACGAGCTTTT | 22.3 | 0.2 | 63.6 | **** |
| *ENO2-GPBP1* | AAAGTGCTGGGATTACAGGG | GATGACGACTCATTTAATTT | 32.3 | 4.9 | 63.6 | **** |
| *ELOVL5-PTP4A1* | GCCGCCCCTTGGGCTAAAAG | CAATTCTGTGGTGTTCTTGG | 22.6 | 0.2 | 63.6 | **** |
| *B2M-UBC* | TGGCCTGGAGGCTATCCAGC | ACAATGCAGATCTTCGTGAA | 24.4 | 1.2 | 61.4 | **** |
| *HNRNPK-UBQLN1* | CACTTGTTCGCGGCCTATAG | TTTAAGGAAGAAATCTCTAA | 22.3 | 0.2 | 61.4 | **** |
| *ZEB1-BMI1* | GAACCCGCGGCGCAATAACG | GATTTTTTATCAAGCAGAAA | 22.1 | 0.2 | 59.2 | **** |
| *OAZ1-RNF126* | GGCGGCAGCAGCAGTGAGAG | GATTATATCTGTCCAAGATG | 24.1 | 1.2 | 58.1 | **** |
| *EPS15L1-KLF2* | TCATCCCCCTCTCCCAGCAG | GTGAGAAGCCCTACCACTGC | 23.1 | 0.7 | 58.1 | **** |
| *OAZ1-SF1* | GGCGGCAGCAGCAGTGAGAG | ACTTCCCAAGTAAGAAGCGG | 21.5 | 0.2 | 57.0 | **** |
| *SPIN1-HNRNPK* | GCAGCCTCGGCGGTCAGCAG | GCTACTGCAGCACTGGGGTG | 21.5 | 0.2 | 57.0 | **** |
| *PTBP1-KLF2* | GCTCTGTGTGCCATGGACGG | GTGAGAAGCCCTACCACTGC | 21.3 | 0.2 | 55.9 | **** |
| *G3BP2-CCNG2* | GCCGCCGCCGTTGGCTGGAG | ATGAAGGATTTGGGGGCAGA | 21.5 | 0.2 | 55.9 | **** |
| *DDX5-HNRNPU* | GACCGCGGCCGGGACCGAGG | GGGACGGCAAAACAGAACAG | 20.8 | 0.2 | 55.4 | **** |
| *MAD1L1-MAFK* | GCGACTGCCTCATCTTCAAG | CTACGAGTTCCAGGGAGCTC | 21.0 | 0.2 | 55.4 | **** |
| *MAEA-CTBP1* | AGGAGTACCCGACCCTCAAG | GCGTCCGACCTCCGATCATG | 26.7 | 3.0 | 54.8 | **** |
| *C21orf33-SUMO3* | CAGGCCGGGAAGCCCATCGG | GAGGGTGTGAAGACAGAGAA | 20.5 | 0.2 | 54.8 | **** |
| *C9orf100-DENND1C* | AGACCAGCCTGGGCAACACA | GGCCAGCCAGTGACCTTCAG | 74.6 | 41.0 | 54.8 | **** |
| *YPEL5-PPP1CB* | GGATCTCACCGCCGCTCAGG | TACGAGGATGTCGTCCAGGA | 19.7 | 0.2 | 53.7 | **** |
| *COTL1-KLHL36* | CTTCATCCAGCAGTGCACAG | GGCTGAAATCTCTTTAATGA | 19.7 | 0.2 | 53.7 | **** |
| *ELOVL5-EEF1A1* | GCCGCCCCTTGGGCTAAAAG | GTGTCGTGAAAACTACCCCT | 20.0 | 0.2 | 53.7 | **** |

| | | | | | | |
|---|---|---|---|---|---|---|
| *SRSF5-EEF1A1* | CCAAAGACCCCGTCCGGTAG | GTGTCGTGAAAACTACCCCT | 20.0 | 0.2 | 53.7 | **** |
| *ZNF292-PNRC1* | ACTGTCAGCAGCTGTGCCAG | GTTTTAAAATCAAAGATGGG | 20.3 | 0.2 | 53.7 | **** |
| *PTBP1-RNF126* | GCTCTGTGTGCCATGGACGG | GATTATATCTGTCCAAGATG | 20.3 | 0.5 | 52.6 | **** |
| *METTL13-ULK2* | GACTACTTTAGATACCTCAT | GCAGTGTCAGCCTACTGTGT | 19.7 | 0.2 | 52.6 | **** |
| *UNKL-UBE2I* | GCGAGCAAGACAGCAAGCAG | GGACTTTGAACATGTCGGGG | 19.5 | 0.2 | 51.8 | **** |
| *SENP3_-ATP6V0C* | TCTGCGAGCCAGGATTCCCG | CCCTGGGCGCTGCCTATGGC | 19.5 | 0.2 | 51.6 | **** |
| *FUS-SETD1A* | CGCGGACATGGCCTCAAACG | TGTAAATGAGCAAAGATGGA | 19.2 | 0.2 | 50.5 | **** |
| *SRGN-KLF6* | TCTGGAATCCTCAGTTCAAG | ACCTGCCTAGAGCTGGAACG | 19.2 | 0.2 | 50.5 | **** |
| *OAZ1-REXO1* | GGCGGCAGCAGCAGTGAGAG | GGCTGGGTTACGACCCCTAC | 19.2 | 0.2 | 50.5 | **** |
| *OAZ1-SCO2* | GGCGGCAGCAGCAGTGAGAG | GAGCATCAGATCCATGCTGC | 26.2 | 3.5 | 49.4 | **** |
| *PIM3-SCO2* | GGTCTACACCGACTTCGACG | GAGCATCAGATCCATGCTGC | 29.2 | 5.2 | 49.4 | **** |
| *B2M-GNAS* | TGGCCTGGAGGCTATCCAGC | GTGCTGGAGAATCTGGTAAA | 44.9 | 15.2 | 49.4 | **** |
| *ACTB-ATP6V0C* | CCGTCCACACCCGCCGCCAG | CCCTGGGCGCTGCCTATGGC | 19.7 | 0.5 | 49.4 | **** |
| *PSPC1-ZMYM2* | GCTTCGGCTTCATCCGCTTG | GACCAAGAATCGCCTTCAGC | 18.7 | 0.2 | 48.3 | **** |
| *USP10-KLHL36* | TGGCCCTCCACAGCCCGCAG | GGCTGAAATCTCTTTAATGA | 18.7 | 0.2 | 48.3 | **** |
| *ANP32E-PLEKHO1* | GGAACAGATCCCCGGAGGAG | GGACCTCAGGATGGAAACCA | 18.7 | 0.2 | 48.3 | **** |
| *HNRNPA2B1-BMI1* | CGACTGAGTCCGCGATGGAG | GATTTTTTATCAAGCAGAAA | 18.2 | 0.2 | 47.2 | **** |
| *SENP3_-SF1* | TCTGCGAGCCAGGATTCCCG | ACTTCCCAAGTAAGAAGCGG | 18.2 | 0.2 | 47.2 | **** |
| *CDKN1A-EEF1A1* | CACCGAGGCACTCAGAGGAG | GTGTCGTGAAAACTACCCCT | 18.5 | 0.2 | 47.2 | **** |
| *LITAF-WDR74* | CTGAGTACCTGGGACCACAG | CACCCTCATCACATGTGTGG | 21.0 | 1.2 | 47.2 | **** |
| *POTEH-RP9* | CCAAAGTGCTAGGATTACAG | GTTGGCGTTGCAAACGCTAT | 21.5 | 1.6 | 47.2 | **** |
| *C6orf115-EEF1A1* | AAGGAAAACCGCGCAGAGAG | GTGTCGTGAAAACTACCCCT | 17.4 | 0.2 | 46.1 | **** |
| *NCOR1-UBB* | CCACGCTTAGCCAGCTCCCG | GTCAAAATGCAGATCTTCGT | 23.8 | 3.0 | 46.1 | **** |
| *RNF126-PTBP1* | AGATCGTCCCGCGCCTGCCG | CGGGGATCTGACGAGCTTTT | 17.9 | 0.2 | 46.1 | **** |
| *POTEH-SMCHD1* | CCAAAGTGCTAGGATTACAG | CCCTCTGATTCTGTTCACAT | 19.0 | 0.9 | 45.5 | **** |
| *USP39-ZFP90* | TTTTCAGATCACCAAACAAG | AGTCTTGCTCTGTTGCCCAG | 23.1 | 2.8 | 45.0 | **** |
| *MBP-CTDP1* | GAGCAACGAGCGCAGCGCCG | AGCGGTTCTGGTGAGGTTGG | 18.2 | 0.7 | 43.9 | **** |
| *B2M-HNRNPUL1* | TGGCCTGGAGGCTATCCAGC | ATGCCATGGACAATATTACC | 21.5 | 2.1 | 43.9 | **** |
| *DDX5-SF1* | GACCGCGGCCGGGACCGAGG | ACTTCCCAAGTAAGAAGCGG | 16.4 | 0.2 | 42.8 | **** |
| *ATP11A-ING1* | GGACGCTCGTGCACAGATAC | AGATCCTGAAGGAGCTAGAC | 18.2 | 0.9 | 42.8 | **** |
| *FOSB-UBC* | CCACCGCCGCCGCCTCCCAG | ACAATGCAGATCTTCGTGAA | 16.7 | 0.2 | 42.8 | **** |
| *PTMA-KLF11* | GCTCCGAAATCACCACCAAG | GTTGACATCATGGACATATG | 16.9 | 0.2 | 42.8 | **** |
| *OAZ1-RPS15* | GGCGGCAGCAGCAGTGAGAG | CGAGCAGCTGATGCAGCTGT | 16.9 | 0.2 | 42.8 | **** |
| *MBP-PQLC1* | GAGCAACGAGCGCAGCGCCG | CATCAAGATGGTGCTCATGT | 21.8 | 2.6 | 42.2 | **** |
| *HNRNPR-C19orf47* | AACCTGGAATTGGAACGGAG | ATGGAGTCTCCCTCTGTTGC | 17.4 | 0.7 | 41.7 | **** |
| *NFATC1-PQLC1* | CGCCGCCGCGGCCGCCCCAG | CATCAAGATGGTGCTCATGT | 17.4 | 0.7 | 41.7 | **** |
| *RNF126-BSG* | AGATCGTCCCGCGCCTGCCG | CCGGCACAGTCTTCACTACC | 20.8 | 2.1 | 41.7 | **** |
| *GLTSCR2-DHX34* | CTACAGGAGCGCACGAGCGG | GGGCTGAGGAAGCTGCCCTC | 23.1 | 3.3 | 41.2 | **** |
| *PIM3-BRD1* | GGTCTACACCGACTTCGACG | GTAATCATTACCAAATGAGG | 16.2 | 0.2 | 35.3 | **** |
| *DDX5-ATP6V0C* | GACCGCGGCCGGGACCGAGG | CCCTGGGCGCTGCCTATGGC | 16.2 | 0.2 | 34.0 | **** |
| *TNRC18-ACTB* | CGTGAACCCAGCCGCGCCAG | GGCGTGATGGTGGGCATGGG | 16.2 | 0.2 | 33.3 | **** |
| *B2M-KLF2* | TGGCCTGGAGGCTATCCAGC | GTGAGAAGCCCTACCACTGC | 16.2 | 0.2 | 32.9 | **** |
| *ADAP1-MAFK* | GCAGGAGCCCTACTCGGCAG | CTACGAGTTCCAGGGAGCTC | 17.7 | 0.9 | 32.7 | **** |
| *YPEL5-UBC* | GGATCTCACCGCCGCTCAGG | ACAATGCAGATCTTCGTGAA | 15.9 | 0.2 | 32.5 | **** |
| *GNB1-SSU72* | GCCGCGACGAGGGCGCTGAG | CAAACGGGGATTCAGCGTCC | 17.2 | 0.9 | 32.4 | **** |
| *B2M-PNRC1* | TGGCCTGGAGGCTATCCAGC | GTTTTAAAATCAAAGATGGG | 15.6 | 0.2 | 32.4 | **** |
| *TPM4-ACTB* | AGCGCGAGCGGCGCGAGAAA | CTCACCATGGATGATGATAT | 16.4 | 0.5 | 32.4 | **** |
| *HP1BP3-CDC42* | AGGACCGGCCGAACGCAGAG | GTCATCATCAGATTTGAAAT | 15.4 | 0.2 | 31.8 | **** |
| *FBRSL1-ANKLE2* | CACTGCAGCCCTCCAAGCAG | GTGAAATGACAATGGATGCT | 15.4 | 0.2 | 27.2 | **** |
| *TMEM91-HNRNPUL1* | GTGAAGCCACGGAAGATAAG | ATGCCATGGACAATATTACC | 15.9 | 0.5 | 26.9 | **** |

| | | | | | | |
|---|---|---|---|---|---|---|
| *EDF1-C9orf86* | CGCAGAAGGACCTGGCCACG | TGAAGATAGTGATCCGGGGA | 15.1 | 0.2 | 26.6 | **** |
| *B2M-YWHAZ* | TGGCCTGGAGGCTATCCAGC | AACATCCAGTCATGGATAAA | 15.1 | 0.2 | 25.6 | **** |
| *PTMA-ATP6V0C* | GCTCCGAAATCACCACCAAG | CCCTGGGCGCTGCCTATGGC | 15.6 | 0.5 | 24.9 | **** |
| *DDX5-HNRNPH3* | GACCGCGGCCGGGACCGAGG | CATTTAAATCAAACGGTATT | 15.6 | 0.5 | 24.8 | **** |
| *ZYX-ACTB* | CGCGCACGCTCCGGCCTGCG | CTCACCATGGATGATGATAT | 15.6 | 0.7 | 23.8 | **** |
| *B2M-CCNG2* | TGGCCTGGAGGCTATCCAGC | ATGAAGGATTTGGGGGCAGA | 15.6 | 0.5 | 23.8 | **** |
| *TTYH3-MAD1L1* | AGGACACCGACTACCAGCAG | GCCACCAGCCCCTCGGGTTC | 15.4 | 0.7 | 23.1 | **** |
| *YPEL5-KLF11* | GGATCTCACCGCCGCTCAGG | GTTGACATCATGGACATATG | 14.1 | 0.2 | 22.4 | **** |
| *OAZ1-MOBKL2A* | GGCGGCAGCAGCAGTGAGAG | GTCCGCGGGAGAGCTGTGGG | 19.0 | 2.3 | 21.7 | **** |
| *PQLC1-CTDP1* | CAGGCGCCGCTCCTTTACAG | AGCGGTTCTGGTGAGGTTGG | 16.9 | 1.4 | 21.6 | **** |
| *ANP32A-CHD2* | GGAACAGGACGCCCTCTGAT | GTGACTTAAGAGATTAAAAT | 15.1 | 0.7 | 21.3 | **** |
| *FUS-ATP6V0C* | CGCGGACATGGCCTCAAACG | CCCTGGGCGCTGCCTATGGC | 13.8 | 0.2 | 20.9 | **** |
| *BANP-SLC7A5* | CACCATCTACGGCATCCGGT | GTGATGTGTCCAATCTAGAT | 13.8 | 0.2 | 20.9 | **** |
| *RASA3-ING1* | GAGCGTGAAGATCAAGATCG | AGATCCTGAAGGAGCTAGAC | 15.4 | 0.9 | 20.5 | **** |
| *PTBP1-KLF16* | GCTCTGTGTGCCATGGACGG | GGGAACGCCCTTTTGCTTGT | 13.6 | 0.2 | 19.8 | **** |
| *ADSS-ZNF238* | ACATCGTGTGCCGCTGCCAG | GTTATGAAGACAGTATGGAG | 13.6 | 0.2 | 19.8 | **** |
| *CEP170-ZNF238* | CCCTCTCTCCACGGTAAGGG | GTTATGAAGACAGTATGGAG | 13.3 | 0.2 | 19.8 | **** |
| *PRPS1-SMCHD1* | CTGAGTAGCTGGGATTACAG | GTTGTTAAAATTACACACTG | 13.3 | 0.2 | 19.6 | **** |
| *OAZ1-METRNL* | GGCGGCAGCAGCAGTGAGAG | CGCCGTGCCGTCCCTGCAGT | 22.3 | 4.4 | 19.3 | **** |
| *OAZ1-ACTB* | GGCGGCAGCAGCAGTGAGAG | GGCGTGATGGTGGGCATGGG | 13.1 | 0.2 | 19.0 | **** |
| *ELMO1-ACTB* | TGTCCTGCGACCCCAAGCAG | CTCACCATGGATGATGATAT | 13.1 | 0.2 | 18.4 | **** |
| *B2M-METRNL* | TGGCCTGGAGGCTATCCAGC | CGGGCTGACGCACGAGGCAC | 15.4 | 1.2 | 18.3 | **** |
| *ARHGEF7-ING1* | GGGCTTCCCTGCGGCTGGAG | AGATCCTGAAGGAGCTAGAC | 15.9 | 1.4 | 18.0 | **** |
| *OAZ1-LPAR2* | GGCGGCAGCAGCAGTGAGAG | ATGGTCATCATGGGCCAGTG | 12.8 | 0.2 | 17.9 | **** |
| *B2M-CDC42* | TGGCCTGGAGGCTATCCAGC | GTCATCATCAGATTTGAAAT | 12.8 | 0.2 | 17.6 | **** |
| *SENP3_-CCNG2* | TCTGCGAGCCAGGATTCCCG | ATGAAGGATTTGGGGGCAGA | 12.8 | 0.2 | 17.3 | **** |
| *DDX3X-UBB* | CGCTCGGGCTGGACCAGCAG | GTCAAAATGCAGATCTTCGT | 12.8 | 0.2 | 17.2 | **** |
| *CPSF6-EEF1A1* | TCGGCGAAGAGTTCAACCAG | GTGTCGTGAAAACTACCCCT | 12.8 | 0.2 | 17.0 | **** |
| *ZEB2-YPEL5* | CTACAAACAAGACTTCGCAG | GTTTTTAGAACTTCAGCCAT | 12.6 | 0.2 | 16.9 | **** |
| *DDX5-CHD2* | GACCGCGGCCGGGACCGAGG | GTGACTTAAGAGATTAAAAT | 12.6 | 0.2 | 16.5 | **** |
| *ACTB-METRNL* | CCGTCCACACCCGCCGCCAG | CGGGCTGACGCACGAGGCAC | 13.1 | 0.5 | 15.7 | **** |
| *ANP32A-B2M* | GGAACAGGACGCCCTCTGAT | GTACTCCAAAGATTCAGGTT | 21.3 | 4.4 | 15.4 | **** |
| *CTBP1-MAEA* | CAACAAGGGCCTGCCGCTTG | TCTCTGCTTTCTATCCGTCA | 12.3 | 0.2 | 14.1 | **** |
| *B2M-NFIL3* | TGGCCTGGAGGCTATCCAGC | ATAACGAAGGAGGCCCAACT | 13.3 | 0.7 | 13.9 | **** |
| *KDM4C-UHRF2* | CGCCCCAACCCGCGCGCCAG | TTGGAAAATGGATATACCTT | 12.1 | 0.2 | 13.6 | **** |
| *ZBTB4-EIF5A* | CGCGGCGCCATCGCAGTCAG | TTGGAATCGAAGCCTCTTAA | 12.1 | 0.2 | 13.0 | **** |
| *HNRNPK-CHD2* | CACTTGTTCGCGGCCTATAG | GTGACTTAAGAGATTAAAAT | 12.1 | 0.2 | 12.9 | **** |
| *NCOR2-UBC* | ACCGCTGCCACCTCCGCGAG | ACAATGCAGATCTTCGTGAA | 47.7 | 23.9 | 12.7 | **** |
| *LOC643201-LOC728554* | CGCCGAGGCTGCTCACAAAA | TCTTCCCACACAGGTTGAGA | 30.5 | 10.5 | 12.7 | **** |
| *HNRNPK-NFIL3* | CACTTGTTCGCGGCCTATAG | ATAACGAAGGAGGCCCAACT | 13.1 | 0.7 | 12.5 | **** |
| *HNRNPU-ZNF238* | GGGCCAGCAGCAGGCGGGAG | GTTATGAAGACAGTATGGAG | 11.8 | 0.2 | 12.4 | **** |
| *FAM60A-GNAS* | GGAACGGCTGCTTTTGGAAG | GTGCTGGAGAATCTGGTAAA | 11.8 | 0.2 | 12.2 | **** |
| *B2M-ACTB* | TGGCCTGGAGGCTATCCAGC | CTCACCATGGATGATGATAT | 11.8 | 0.2 | 12.2 | **** |
| *SENP3_-OAZ1* | TCTGCGAGCCAGGATTCCCG | GATGATCGGCTGAATGTAAC | 11.8 | 0.2 | 12.1 | **** |
| *SLC22A20-WDR33* | GCAGCCACCCCATCTGGGAA | TATCCTAGAAGGAAAACATC | 14.6 | 1.4 | 12.1 | **** |
| *LOC100289019-DEDD2* | TGAGAAACCCGGCAATGAAA | GCTGGAGTGCAATGGCACAA | 24.1 | 6.6 | 11.0 | **** |
| *KLF2-ACTB* | GCGAGCGCGGCCTGCAGGAG | CTCACCATGGATGATGATAT | 12.8 | 0.7 | 10.5 | **** |
| *YPEL5-GNAS* | GGATCTCACCGCCGCTCAGG | GTGCTGGAGAATCTGGTAAA | 11.5 | 0.2 | 10.3 | **** |
| *YPEL5-EEF1A1* | GGATCTCACCGCCGCTCAGG | GTGTCGTGAAAACTACCCCT | 11.5 | 0.2 | 9.9 | **** |
| *PTMA-CHD2* | GCTCCGAAATCACCACCAAG | GTGACTTAAGAGATTAAAAT | 11.5 | 0.2 | 9.4 | **** |

| | | | | | | |
|---|---|---|---|---|---|---|
| *SMOX-LOC389906* | CCAAAGTGCTGGAATTACAG | GTTGAAAAAGATCATCTAGC | 16.4 | 2.3 | 9.3 | **** |
| *OAZ1-UBB* | GGCGGCAGCAGCAGTGAGAG | GTCAAAATGCAGATCTTCGT | 15.4 | 1.9 | 9.2 | **** |
| *CSNK1G2-BTBD2* | CCGGCCACGGCGGAGCCAAG | GTTCGTGCTGGCCGTGGGCA | 13.1 | 0.9 | 8.9 | **** |
| *YWHAZ-EEF1A1* | ACCACCCACTCCGGACACAG | GTGTCGTGAAAACTACCCCT | 11.3 | 0.2 | 8.9 | **** |
| *DDX5-GNAS* | GACCGCGGCCGGGACCGAGG | GTGCTGGAGAATCTGGTAAA | 11.3 | 0.2 | 7.8 | **** |
| *FAM60A-CHD2* | GGAACGGCTGCTTTTGGAAG | GTGACTTAAGAGATTAAAAT | 11.3 | 0.2 | 7.5 | **** |
| *ELL-KLF2* | AGAGCTACCGCGCCAGACAG | GTGAGAAGCCCTACCACTGC | 11.3 | 0.2 | 7.5 | **** |
| *ZEB1-GNAS* | GAACCCGCGGCGCAATAACG | GTGCTGGAGAATCTGGTAAA | 12.8 | 0.9 | 7.2 | **** |
| *ACTB-SCO2* | CCGTCCACACCCGCCGCCAG | GAGCATCAGATCCATGCTGC | 12.3 | 0.7 | 7.2 | **** |
| *TTYH3-LFNG* | AGGACACCGACTACCAGCAG | ACGTTCATCTTCACTGACGG | 12.8 | 0.9 | 7.1 | **** |
| *USP12-ZMYM2* | TCGCCTCCATCTGTACCATG | GACCAAGAATCGCCTTCAGC | 11.0 | 0.2 | 6.8 | **** |
| *TAF9B-VPS13B* | TTTCTGCATTTCCAACTGAG | GATGACCATGAAAGCTGTGG | 11.0 | 0.2 | 6.4 | **** |
| *N4BP2L1-ZMYM2* | GGGAAAACTACACTGGCCAG | GACCAAGAATCGCCTTCAGC | 11.0 | 0.2 | 6.0 | **** |
| *SS18L1-RBM38* | CGCAGCAAACCATCCAGAAG | GCTGACCCCGCACTACATCT | 12.6 | 0.9 | 6.0 | **** |
| *FUS-ACTB* | CGCGGACATGGCCTCAAACG | CTCACCATGGATGATGATAT | 10.8 | 0.2 | 6.0 | **** |
| *UBE2L3-SIAH2* | AGACCATCCTGGCTAACACG | TATGCCACCACGGGCTGTTC | 40.8 | 19.4 | 5.8 | **** |
| *LFNG-ACTB* | TCTCGCGCCACAAGGAGATG | CTCACCATGGATGATGATAT | 10.5 | 0.2 | 5.6 | **** |
| *TPM4-METRNL* | AGCGCGAGCGGCGCGAGAAA | CGGGCTGACGCACGAGGCAC | 10.5 | 0.2 | 5.6 | **** |
| *USP6-L3MBTL4* | CCCACTGATTCTACATTATG | CAGATCGATGGCAAAGCCTT | 10.3 | 0.2 | 5.4 | **** |
| *MAFK-ACTB* | CGCCCTCCGCGCGACGCCAG | CTCACCATGGATGATGATAT | 10.3 | 0.2 | 5.1 | **** |
| *CSNK1G2-KLF16* | CCGGCCACGGCGGAGCCAAG | GGGAACGCCCTTTTGCTTGT | 10.3 | 0.2 | 5.0 | **** |
| *PTBP1-ACTB* | GCTCTGTGTGCCATGGACGG | CTCACCATGGATGATGATAT | 10.3 | 0.2 | 4.7 | **** |
| *CPSF6-DDX5* | TCGGCGAAGAGTTCAACCAG | GTTTGGTGCACCTCGATTTG | 10.3 | 0.2 | 4.0 | **** |
| *PIK3R5-UBB* | GCGCCCGGCGGGGACTGCAG | GTCAAAATGCAGATCTTCGT | 10.0 | 0.2 | 3.8 | **** |
| *ZFPM1-SLC7A5* | CAGTCCCTGGAGCGGGCCAG | GTGATGTGTCCAATCTAGAT | 10.0 | 0.2 | 3.7 | **** |
| *TRNAN35-CHD2* | TGAGGACTGCCTGTTTGCAG | GTGACTTAAGAGATTAAAAT | 10.0 | 0.2 | 3.5 | **** |
| *LOC100129917-CTBP1* | GCACCTTCCCGCCTTCTCAG | TAAGACTTTGACCCCGACGC | 11.5 | 1.2 | 2.7 | **** |
| *MANBA-UBE2D3* | TCCAGCAGGGCCTGATCCAG | GAACTTAGTGATTTGGCCCG | 26.4 | 10.1 | 2.4 | **** |
| *CSNK1D-METRNL* | CTTCGGAGACATCTATCTCG | CGGGCTGACGCACGAGGCAC | 12.8 | 1.9 | 1.9 | **** |
| *CPD-TJP2* | ACATTGGTATGATGTGGAAG | AGTCTTGCTGTGTTGCCCAG | 37.4 | 18.7 | 1.9 | **** |
| *TRABD-SCO2* | AGCGCCGCTGCCTCGCCTCG | GAGCATCAGATCCATGCTGC | 12.8 | 2.1 | 1.9 | **** |
| *CSNK1G2-MOBKL2A* | CCGGCCACGGCGGAGCCAAG | GTCCGCGGGAGAGCTGTGGG | 10.3 | 0.9 | 1.7 | **** |
| *OAZ1-RNF166* | GGCGGCAGCAGCAGTGAGAG | GTTCTGCGGGGAGTGTCTCC | 10.8 | 1.6 | 1.6 | **** |
| *CTDP1-PQLC1* | GCAGGGGGCCGGGGGCCCAG | CATCAAGATGGTGCTCATGT | 12.1 | 2.6 | 1.2 | **** |
| *SLC16A3-METRNL* | AGGCGGCCCAGCGGCGGCAG | CGGGCTGACGCACGAGGCAC | 19.7 | 7.3 | 1.1 | **** |
| *AMZ2-AMZ2* | CTACCTACAGGACCTTAAAG | GTGTGGGGATATTCAGCTTT | 31.3 | 15.9 | 1.0 | **** |
| *KLK10-C17orf87* | AAAGTGCTGGGATTACAAGC | CCCCACAGGAAGCCCCAAGT | 12.6 | 3.7 | 1.0 | **** |
| *ARF1-H3F3A* | TCGGAGCAGCAGCCTCTGAG | GTAAGTAAGGAGGTCTCTGT | 12.3 | 4.2 | 1.0 | **** |
| *RRN3P1-RRN3P2* | TTCTAAGAAAAGCTTTTCAG | GACACCGTGGTTTCTCATGC | 18.5 | 8.4 | 0.8 | **** |
| *FAM126B-EIF6* | AATGGTTATCAGAATTCAAG | ATGGAGTCTCGCTCTGTCAC | 26.2 | 18.3 | 0.8 | ** |
| *DLEU2-C20orf152* | TTCCTTCTACTGATCTCCAA | ACAGAGTCTTGCTTTGTCAC | 11.5 | 6.3 | 0.4 | ** |
| *RB1-ITM2B* | CATCTGTGGATGGAGTATTG | GACCCAGATGATGTGGTACC | 20.8 | 20.4 | 0.0 | |
| *KIAA1267-ARL17A* | AACAGATACGTGCTAATAAG | GTTTCTGTGTGGAGACAGTA | 36.4 | 38.4 | -0.1 | |
| *SQRDL-B2M* | AAGGCTCCGCTCACGCCCGG | GTACTCCAAAGATTCAGGTT | 20.3 | 27.4 | -0.3 | ** |
| | | Averages | 23.901024 | 2.12315 | | |

** $p$ <0.005; **** $p$ <0.0002

**Table S2 5'-*OAZ1*-HFGs encoded diversities of putative proteins**

| HFG IDs | 5' Junction Sequences | 3' Junction Sequences | RFs (%) | Protein Alterations |
|---|---|---|---|---|
| *OAZ1-ACTB* | GGCGGCAGCAGCAGTGAGAG | GGCGTGATGGTGGGCATGGG | 13.1 | truncated ACTB |
| *OAZ1-ATP6V0C* | GGCGGCAGCAGCAGTGAGAG | CCCTGGGCGCTGCCTATGGC | 33.6 | truncated ATP6V0C |
| *OAZ1-BTBD2* | GGCGGCAGCAGCAGTGAGAG | GTTCGTGCTGGCCGTGGGCA | 39.2 | OAZ1-BTBD2 Hybrid |
| *OAZ1-CCNG2* | GGCGGCAGCAGCAGTGAGAG | ATGAAGGATTTGGGGGCAGA | 24.4 | truncated CCNG2 |
| *OAZ1-CSNK1G2* | GGCGGCAGCAGCAGTGAGAG | CTGTGAGCCGTGAGCTTTGA | 30 | Intact CSNK1G2 |
| *OAZ1-DAZAP1* | GGCGGCAGCAGCAGTGAGAG | GAAGCTCTTCGTGGGCGGTC | 26.4 | OAZ1-DAZAP1 Hybrid |
| *OAZ1-DDX5* | GGCGGCAGCAGCAGTGAGAG | GTTTGGTGCACCTCGATTTG | 23.1 | OAZ1-DDX5 Hybrid |
| *OAZ1-DOT1L* | GGCGGCAGCAGCAGTGAGAG | GTAACTAGGATTTCTACCTC | 36.9 | Very Short OAZ1 |
| *OAZ1-KLF16* | GGCGGCAGCAGCAGTGAGAG | GGGAACGCCCTTTTGCTTGT | 67.9 | Very Short OAZ1 |
| *OAZ1-KLF2* | GGCGGCAGCAGCAGTGAGAG | GTGAGAAGCCCTACCACTGC | 47.9 | Very Short OAZ1 |
| *OAZ1-LIMD2* | GGCGGCAGCAGCAGTGAGAG | AACCCAGCGGGTGCCGCTTC | 25.6 | Intact LIMD2 |
| *OAZ1-LPAR2* | GGCGGCAGCAGCAGTGAGAG | ATGGTCATCATGGGCCAGTG | 12.8 | OAZ1-LPAR2 Hybrid |
| *OAZ1-LRCH4* | GGCGGCAGCAGCAGTGAGAG | CCCACAGCCCCCATCCCGTA | 31.5 | OAZ1-LRCH4 Hybrid |
| *OAZ1-METRNL* | GGCGGCAGCAGCAGTGAGAG | CGCCGTGCCGTCCCTGCAGT | 22.3 | Very Short OAZ1 |
| *OAZ1-MOBKL2A* | GGCGGCAGCAGCAGTGAGAG | GTCCGCGGGAGAGCTGTGGG | 19.0 | Intact MOBKL2A |
| *OAZ1-PTBP1* | GGCGGCAGCAGCAGTGAGAG | CGGGGATCTGACGAGCTTTT | 22.3 | truncated PTBP1 |
| *OAZ1-REXO1* | GGCGGCAGCAGCAGTGAGAG | GGCTGGGTTACGACCCCTAC | 19.2 | truncated REXO1 |
| *OAZ1-RNF126* | GGCGGCAGCAGCAGTGAGAG | GATTATATCTGTCCAAGATG | 24.1 | truncated RNF126 |
| *OAZ1-RNF166* | GGCGGCAGCAGCAGTGAGAG | GTTCTGCGGGGAGTGTCTCC | 10.8 | OAZ1-RNF166 Hybrid |
| *OAZ1-RPS15* | GGCGGCAGCAGCAGTGAGAG | CGAGCAGCTGATGCAGCTGT | 16.9 | OAZ1-RPS15 Hybrid |
| *OAZ1-SCO2* | GGCGGCAGCAGCAGTGAGAG | GAGCATCAGATCCATGCTGC | 26.2 | OAZ1-SCO2 Hybrid |
| *OAZ1-SF1* | GGCGGCAGCAGCAGTGAGAG | ACTTCCCAAGTAAGAAGCGG | 21.5 | truncated SF1 |
| *OAZ1-UBB* | GGCGGCAGCAGCAGTGAGAG | GTCAAAATGCAGATCTTCGT | 15.4 | Intact UBB |
| *OAZ1-ZBTB7A* | GGCGGCAGCAGCAGTGAGAG | GTCTCGGCGCGGAAGATGGC | 29.2 | Intact ZBTB7A |

RF: recurrent frequency;

Fig.S1

a). ACTB-ATP6V0C

```
ATP6V0C       RRETGPTLRRSRGRRRPETPAEAEAGRGRAPGPGRWAGGTARGRSTGRPGVPGTRLPHGAK
ACTB-ATP6V0C  -------------------------------------------------------------

ATP6V0C       WYGSQGRGPGHVTRPRPPFCSAVLVFRAQRLTGRIAFAAARPQTFVPGPSSPPPPPPPRPA
ACTB-ATP6V0C  -------------------------------------------------------------

ATP6V0C       ELAPSPPADMSESKSGPEYASFFAVMGASAAMVFSALGAAYGTAKSGTGIAAMSVMRPEQI
ACTB-ATP6V0C  ----------------------------------------------------MSVMRPEQI

ATP6V0C       MKSIIPVVMAGIIAIYGLVVAVLIANSLNDDISLYKSFLQLGAGLSVGLSGLAAGFAIGIV
ACTB-ATP6V0C  MKSIIPVVMAGIIAIYGLVVAVLIANSLNDDISLYKSFLQLGAGLSVGLSGLAAGFAIGIV

ATP6V0C       GDAGVRGTAQQPRLFVGMILILIFAEVLGLYGLIVALILSTK
ACTB-ATP6V0C  GDAGVRGTAQQPRLFVGMILILIFAEVLGLYGLIVALILSTK
```

b). ACTB-EEF1A1

```
EEF1A1       MGKEKTHINIVVIGHVDSGKSTTTGHLIYKCGGIDKRTIEKFEKEAAEMGKGSFKYAWVLD
ACTB-EEF1A1  MGKEKTHINIVVIGHVDSGKSTTTGHLIYKCGGIDKRTIEKFEKEAAEMGKGSFKYAWVLD

EEF1A1       KLKAERERGITIDISLWKFETSKYYVTIIDAPGHRDFIKNMITGTSQADCAVLIVAAGVGE
ACTB-EEF1A1  KLKAERERGITIDISLWKFETSKYYVTIIDAPGHRDFIKNMITGTSQADCAVLIVAAGVGE

EEF1A1       FEAGISKNGQTREHALLAYTLGVKQLIVGVNKMDSTEPPYSQKRYEEIVKEVSTYIKKIGY
ACTB-EEF1A1  FEAGISKNGQTREHALLAYTLGVKQLIVGVNKMDSTEPPYSQKRYEEIVKEVSTYIKKIGY

EEF1A1       NPDTVAFVPISGWNGDNMLEPSANMPWFKGWKVTRKDGNASGTTLLEALDCILPPTRPTDK
ACTB-EEF1A1  NPDTVAFVPISGWNGDNMLEPSANMPWFKGWKVTRKDGNASGTTLLEALDCILPPTRPTDK

EEF1A1       PLRLPLQDVYKIGGIGTVPVGRVETGVLKPGMVVTFAPVNVTTEVKSVEMHHEALSEALPG
ACTB-EEF1A1  PLRLPLQDVYKIGGIGTVPVGRVETGVLKPGMVVTFAPVNVTTEVKSVEMHHEALSEALPG

EEF1A1       DNVGFNVKNVSVKDVRRGNVAGDSKNDPPMEAAGFTAQVIILNHPGQISAGYAPVLDCHTA
ACTB-EEF1A1  DNVGFNVKNVSVKDVRRGNVAGDSKNDPPMEAAGFTAQVIILNHPGQISAGYAPVLDCHTA

EEF1A1       HIACKFAELKEKIDRRSGKKLEDGPKFLKSGDAAIVDMVPGKPMCVESFSDYPPLGRFAVR
ACTB-EEF1A1  HIACKFAELKEKIDRRSGKKLEDGPKFLKSGDAAIVDMVPGKPMCVESFSDYPPLGRFAVR
```

```
EEF1A1        DMRQTVAVGVIKAVDKKAAGAGKVTKSAQKAQKAK-
ACTB-EEF1A1   DMRQTVAVGVIKAVDKKAAGAGKVTKSAQKAQKAK-
```

c). ACTB-KLF2

```
KLF2         MALSEPILPSFSTFASPCRERGLQERWPRAEPESGGTDDDLNSVLDFILSMGLDGLGAEAA
ACTB-KLF2    -------------------------------------------------------------

KLF2         PEPPPPPPPPPAFYYPEPGAPPPYSAPAGGLVSELLRPELDAPLGPALHGRFLLAPPGRLVK
ACTB-KLF2    --------------------------------------------------------------

KLF2         AEPPEADGGGGYGCAPGLTRGPRGLKREGAPGPAASCMRGPGGRPPPPPDTPPLSPDGPAR
ACTB-KLF2    -------------------------------------------------------------

KLF2         LPAPGPRASFPPPFGGPGFGAPGPGLHYAPPAPPAFGLFDDAAAAAAALGLAPPAARGLLT
ACTB-KLF2    -------------------------------------------------------------

KLF2         PPASPLELLEAKPKRGRRSWPRKRTATHTCSYAGCGKTYTKSSHLKAHLRTHTGEKPYHCN
ACTB-KLF2    -------LWLERPRR-RPIKPSGATRHHRRDRVRPASTEPRLCRSAARPHPPPGEKPYHCN

KLF2         WDGCGWKFARSDELTRHYRKHTGHRPFQCHLCDRAFSRSDHLALHMKRHM
ACTB-KLF2    WDGCGWKFARSDELTRHYRKHTGHRPFQCHLCDRAFSRSDHLALHMKRHM
```

d). ACTB-MAFK

```
MAFK         MTTNPKPNKALKVKKEAGENAPVLSDDELVSMSVRELNQHLRGLTKEEVTRLKQRRRTLKN
ACTB-MAFK    MTTNPKPNKALKVKKEAGENAPVLSDDELVSMSVRELNQHLRGLTKEEVTRLKQRRRTLKN

MAFK         RGYAASCRIKRVTQKEELERQRVELQQEVEKLARENSSMRLELDALRSKYEALQTFARTVA
ACTB-MAFK    RGYAASCRIKRVTQKEELERQRVELQQEVEKLARENSSMRLELDALRSKYEALQTFARTVA

MAFK         RGPVAPSKVATTSVITIVKSTELSSTSVPFSAAS
ACTB-MAFK    RGPVAPSKVATTSVITIVKSTELSSTSVPFSAAS
```

e). ACTB-METRNL

```
METRNL       M-RGAARAAWGRAGQPWPRPPAPGPP----PPPLPLLLLLLAGLLGGAGAQYSSDRCSWKG
ACTB-METRNL  MARAAAAAPYKTQRRDAPPPPRPRPPREHRASPLPI-----------------RRPSTPAA

METRNL       SGLTHEAHRKEVEQVYLRCAAGAVEWMYPTGALIVNLRPNTFSPARHLTVCIRSFTDSSGA
ACTB-METRNL  SGLTHEAHRKEVEQVYLRCAAGAVEWMYPTGALIVNLRPNTFSPARHLTVCIRSFTDSSGA

METRNL       NIYLEKTGELRLLVPDGDGRPGRVQCFGLEQGGLFVEATPQQDIGRRTTGFQYELVRRHRA
ACTB-METRNL  NIYLEKTGELRLLVPDGDGRPGRVQCFGLEQGGLFVEATPQQDIGRRTTGFQYELVRRHRA

METRNL       SDLHELSAPCRPCSDTEVLLAVCTSDFAVRGSIQQVTHEPERQDSAIHLRVSRLYRQKSRV
ACTB-METRNL  SDLHELSAPCRPCSDTEVLLAVCTSDFAVRGSIQQVTHEPERQDSAIHLRVSRLYRQKSRV

METRNL       FEPVPEGDGHWQGRVRTLLECGVRPGHGDFLFTGHMHFGEARLGCAPRFKDFQRMYRDAQE
ACTB-METRNL  FEPVPEGDGHWQGRVRTLLECGVRPGHGDFLFTGHMHFGEARLGCAPRFKDFQRMYRDAQE

METRNL       RGLNPCEVGTD
ACTB-METRNL  RGLNPCEVGTD
```

f). ACTB-SCO2

```
SCO2       -------------------------------------------------MLLLTRSPTAWH
ACTB-SCO2  MARAAAAAPYKTQRRDAPPPPRPRPPREHRASPLPIRRPSTPAARSIRSMLLLTRSPTAWH

SCO2       RLSQLKPRVLPGTLGGQALHLRSWLLSRQGPAETGGQGQPQGPGLRTRLLITGLFGAGLGG
ACTB-SCO2  RLSQLKPRVLPGTLGGQALHLRSWLLSRQGPAETGGQGQPQGPGLRTRLLITGLFGAGLGG

SCO2       AWLALRAEKERLQQQKRTEALRQAAVGQGDFHLLDHRGRARCKADFRGQWVLMYFGFTHCP
ACTB-SCO2  AWLALRAEKERLQQQKRTEALRQAAVGQGDFHLLDHRGRARCKADFRGQWVLMYFGFTHCP

SCO2       DICPDELEKLVQVVRQLEAEPGLPPVQPVFITVDP
ACTB-SCO2  DICPDELEKLVQVVRQLEAEPGLPPVQPVFITVDP
```

Fig.S1 Analysis of putative proteins generated by 5'-ACTB-fused HFGs.
a). ACTB-ATP6V0C encoded a truncated ATPase H+ transporting V0 subunit c;
b) ACTB-EEF1A1 coded for intact eukaryotic translation elongation factor 1 alpha 1; c). ACTB-KLF2 encoded an ORF without starting condon;
d). ACTB-MAFK coded for intact MAF bZIP transcription factor K;
e). ACTB-METRNL encoded a beta-actin-meteorin like, glial cell differentiation regulator hybrid protein; f). ACTB-SCO2 coded for a beta-actin-synthesis of cytochrome C oxidase 2 hybrid protein.

```
ACTB          -------------------------------------------------------------
B2M-ACTB      -------------------------------------------------------------
FUS-ACTB      -------------------------------------------------------------
MAFK-ACTB     -------------------------------------------------------------
PTBP1-ACTB    -------------------------------------------------------------
ZYX-ACTB      -------------------------------------------------------------
TNRC18-ACTB   -------------------------------------------------------------
OAZ1-ACTB     -------------------------------------------------------------
PTMA-ACTB     -------------------------------------------------------------
ELMO1-ACTB    -------------------------------------------------------------
KLF2-ACTB     -------------------------------------------------------------
TPM4-ACTB     -------------------------------------------------------------
LFNG-ACTB     MLKRCGRRLLLALAGALLACLLVLTADPPPPPLPAERGRRALRSLAGPAGAAPAPGLGAAA

ACTB          -------------------------------------------------------------
B2M-ACTB      -------------------------------------------------------------
FUS-ACTB      -------------------------------------------------------------
MAFK-ACTB     -------------------------------------------------------------
PTBP1-ACTB    -------------------------------------------------------------
ZYX-ACTB      -------------------------------------------------------------
TNRC18-ACTB   -------------------------------------------------------------
OAZ1-ACTB     -------------------------------------------------------------
PTMA-ACTB     ---------------------------------MSDAA-----------------------
ELMO1-ACTB    ---------------------------------MVAGR-----------------------
KLF2-ACTB     -------------------------------MALSEPI-----------------------
TPM4-ACTB     -------------------------------MAGLNSLEAVKRKIQALQQQADEAEDRAQG
LFNG-ACTB     AAPGALVRDVHSLSEYFSLLTRARRDAGPPPGAAPRPADGHPRPLAEPLAPRDVFIAVKTT

ACTB          ------------------------MDDDIAALVVDNGSGMCKAGFAGDDAPRAVFPSIVGR
B2M-ACTB      ------------------------MDDDIAALVVDNGSGMCKAGFAGDDAPRAVFPSIVGR
FUS-ACTB      ------------------------MDDDIAALVVDNGSGMCKAGFAGDDAPRAVFPSIVGR
MAFK-ACTB     ------------------------MDDDIAALVVDNGSGMCKAGFAGDDAPRAVFPSIVGR
PTBP1-ACTB    ------------------------MDDDIAALVVDNGSGMCKAGFAGDDAPRAVFPSIVGR
ZYX-ACTB      ------------------------MDDDIAALVVDNGSGMCKAGFAGDDAPRAVFPSIVGR
TNRC18-ACTB   -------------------------------------------------------------
OAZ1-ACTB     -------------------------------------------------------------
PTMA-ACTB     ----VDTSSEITT--------KLTMDDDIAALVVDNGSGMCKAGFAGDDAPRAVFPSIVGR
ELMO1-ACTB    ----RSSDRREPLPLLSCDPKQLTMDDDIAALVVDNGSGMCKAGFAGDDAPRAVFPSIVGR
KLF2-ACTB     ----LPSFSTFASPCRERGLQELTMDDDIAALVVDNGSGMCKAGFAGDDAPRAVFPSIVGR
TPM4-ACTB     LQRELDGERERRE--------KLTMDDDIAALVVDNGSGMCKAGFAGDDAPRAVFPSIVGR
LFNG-ACTB     KKFHRARLDLLLETWISRHKEMLTMDDDIAALVVDNGSGMCKAGFAGDDAPRAVFPSIVGR

ACTB          PRHQGVMVGMGQKDSYVGDEAQSKRGILTLKYPIEHGIVTNWDDMEKIWHHTFYNELRVAP
B2M-ACTB      PRHQGVMVGMGQKDSYVGDEAQSKRGILTLKYPIEHGIVTNWDDMEKIWHHTFYNELRVAP
FUS-ACTB      PRHQGVMVGMGQKDSYVGDEAQSKRGILTLKYPIEHGIVTNWDDMEKIWHHTFYNELRVAP
MAFK-ACTB     PRHQGVMVGMGQKDSYVGDEAQSKRGILTLKYPIEHGIVTNWDDMEKIWHHTFYNELRVAP
PTBP1-ACTB    PRHQGVMVGMGQKDSYVGDEAQSKRGILTLKYPIEHGIVTNWDDMEKIWHHTFYNELRVAP
ZYX-ACTB      PRHQGVMVGMGQKDSYVGDEAQSKRGILTLKYPIEHGIVTNWDDMEKIWHHTFYNELRVAP
TNRC18-ACTB   ------MVGMGQKDSYVGDEAQSKRGILTLKYPIEHGIVTNWDDMEKIWHHTFYNELRVAP
```

```
OAZ1-ACTB     ------MVGMGQKDSYVGDEAQSKRGILTLKYPIEHGIVTNWDDMEKIWHHTFYNELRVAP
PTMA-ACTB     PRHQGVMVGMGQKDSYVGDEAQSKRGILTLKYPIEHGIVTNWDDMEKIWHHTFYNELRVAP
ELMO1-ACTB    PRHQGVMVGMGQKDSYVGDEAQSKRGILTLKYPIEHGIVTNWDDMEKIWHHTFYNELRVAP
KLF2-ACTB     PRHQGVMVGMGQKDSYVGDEAQSKRGILTLKYPIEHGIVTNWDDMEKIWHHTFYNELRVAP
TPM4-ACTB     PRHQGVMVGMGQKDSYVGDEAQSKRGILTLKYPIEHGIVTNWDDMEKIWHHTFYNELRVAP
LFNG-ACTB     PRHQGVMVGMGQKDSYVGDEAQSKRGILTLKYPIEHGIVTNWDDMEKIWHHTFYNELRVAP

ACTB          EEHPVLLTEAPLNPKANREKMTQIMFETFNTPAMYVAIQAVLSLYASGRTTGIVMDSGDGV
B2M-ACTB      EEHPVLLTEAPLNPKANREKMTQIMFETFNTPAMYVAIQAVLSLYASGRTTGIVMDSGDGV
FUS-ACTB      EEHPVLLTEAPLNPKANREKMTQIMFETFNTPAMYVAIQAVLSLYASGRTTGIVMDSGDGV
MAFK-ACTB     EEHPVLLTEAPLNPKANREKMTQIMFETFNTPAMYVAIQAVLSLYASGRTTGIVMDSGDGV
PTBP1-ACTB    EEHPVLLTEAPLNPKANREKMTQIMFETFNTPAMYVAIQAVLSLYASGRTTGIVMDSGDGV
ZYX-ACTB      EEHPVLLTEAPLNPKANREKMTQIMFETFNTPAMYVAIQAVLSLYASGRTTGIVMDSGDGV
TNRC18-ACTB   EEHPVLLTEAPLNPKANREKMTQIMFETFNTPAMYVAIQAVLSLYASGRTTGIVMDSGDGV
OAZ1-ACTB     EEHPVLLTEAPLNPKANREKMTQIMFETFNTPAMYVAIQAVLSLYASGRTTGIVMDSGDGV
PTMA-ACTB     EEHPVLLTEAPLNPKANREKMTQIMFETFNTPAMYVAIQAVLSLYASGRTTGIVMDSGDGV
ELMO1-ACTB    EEHPVLLTEAPLNPKANREKMTQIMFETFNTPAMYVAIQAVLSLYASGRTTGIVMDSGDGV
KLF2-ACTB     EEHPVLLTEAPLNPKANREKMTQIMFETFNTPAMYVAIQAVLSLYASGRTTGIVMDSGDGV
TPM4-ACTB     EEHPVLLTEAPLNPKANREKMTQIMFETFNTPAMYVAIQAVLSLYASGRTTGIVMDSGDGV
LFNG-ACTB     EEHPVLLTEAPLNPKANREKMTQIMFETFNTPAMYVAIQAVLSLYASGRTTGIVMDSGDGV

ACTB          THTVPIYEGYALPHAILRLDLAGRDLTDYLMKILTERGYSFTTTAEREIVRDIKEKLCYVA
B2M-ACTB      THTVPIYEGYALPHAILRLDLAGRDLTDYLMKILTERGYSFTTTAEREIVRDIKEKLCYVA
FUS-ACTB      THTVPIYEGYALPHAILRLDLAGRDLTDYLMKILTERGYSFTTTAEREIVRDIKEKLCYVA
MAFK-ACTB     THTVPIYEGYALPHAILRLDLAGRDLTDYLMKILTERGYSFTTTAEREIVRDIKEKLCYVA
PTBP1-ACTB    THTVPIYEGYALPHAILRLDLAGRDLTDYLMKILTERGYSFTTTAEREIVRDIKEKLCYVA
ZYX-ACTB      THTVPIYEGYALPHAILRLDLAGRDLTDYLMKILTERGYSFTTTAEREIVRDIKEKLCYVA
TNRC18-ACTB   THTVPIYEGYALPHAILRLDLAGRDLTDYLMKILTERGYSFTTTAEREIVRDIKEKLCYVA
OAZ1-ACTB     THTVPIYEGYALPHAILRLDLAGRDLTDYLMKILTERGYSFTTTAEREIVRDIKEKLCYVA
PTMA-ACTB     THTVPIYEGYALPHAILRLDLAGRDLTDYLMKILTERGYSFTTTAEREIVRDIKEKLCYVA
ELMO1-ACTB    THTVPIYEGYALPHAILRLDLAGRDLTDYLMKILTERGYSFTTTAEREIVRDIKEKLCYVA
KLF2-ACTB     THTVPIYEGYALPHAILRLDLAGRDLTDYLMKILTERGYSFTTTAEREIVRDIKEKLCYVA
TPM4-ACTB     THTVPIYEGYALPHAILRLDLAGRDLTDYLMKILTERGYSFTTTAEREIVRDIKEKLCYVA
LFNG-ACTB     THTVPIYEGYALPHAILRLDLAGRDLTDYLMKILTERGYSFTTTAEREIVRDIKEKLCYVA

ACTB          LDFEQEMATAASSSSLEKSYELPDGQVITIGNERFRCPEALFQPSFLGMESCGIHETTFNS
B2M-ACTB      LDFEQEMATAASSSSLEKSYELPDGQVITIGNERFRCPEALFQPSFLGMESCGIHETTFNS
FUS-ACTB      LDFEQEMATAASSSSLEKSYELPDGQVITIGNERFRCPEALFQPSFLGMESCGIHETTFNS
MAFK-ACTB     LDFEQEMATAASSSSLEKSYELPDGQVITIGNERFRCPEALFQPSFLGMESCGIHETTFNS
PTBP1-ACTB    LDFEQEMATAASSSSLEKSYELPDGQVITIGNERFRCPEALFQPSFLGMESCGIHETTFNS
ZYX-ACTB      LDFEQEMATAASSSSLEKSYELPDGQVITIGNERFRCPEALFQPSFLGMESCGIHETTFNS
TNRC18-ACTB   LDFEQEMATAASSSSLEKSYELPDGQVITIGNERFRCPEALFQPSFLGMESCGIHETTFNS
OAZ1-ACTB     LDFEQEMATAASSSSLEKSYELPDGQVITIGNERFRCPEALFQPSFLGMESCGIHETTFNS
PTMA-ACTB     LDFEQEMATAASSSSLEKSYELPDGQVITIGNERFRCPEALFQPSFLGMESCGIHETTFNS
ELMO1-ACTB    LDFEQEMATAASSSSLEKSYELPDGQVITIGNERFRCPEALFQPSFLGMESCGIHETTFNS
KLF2-ACTB     LDFEQEMATAASSSSLEKSYELPDGQVITIGNERFRCPEALFQPSFLGMESCGIHETTFNS
TPM4-ACTB     LDFEQEMATAASSSSLEKSYELPDGQVITIGNERFRCPEALFQPSFLGMESCGIHETTFNS
LFNG-ACTB     LDFEQEMATAASSSSLEKSYELPDGQVITIGNERFRCPEALFQPSFLGMESCGIHETTFNS

ACTB          IMKCDVDIRKDLYANTVLSGGTTMYPGIADRMQKEITALAPSTMKIKIIAPPERKYSVWIG
```

```
B2M-ACTB     IMKCDVDIRKDLYANTVLSGGTTMYPGIADRMQKEITALAPSTMKIKIIAPPERKYSVWIG
FUS-ACTB     IMKCDVDIRKDLYANTVLSGGTTMYPGIADRMQKEITALAPSTMKIKIIAPPERKYSVWIG
MAFK-ACTB    IMKCDVDIRKDLYANTVLSGGTTMYPGIADRMQKEITALAPSTMKIKIIAPPERKYSVWIG
PTBP1-ACTB   IMKCDVDIRKDLYANTVLSGGTTMYPGIADRMQKEITALAPSTMKIKIIAPPERKYSVWIG
ZYX-ACTB     IMKCDVDIRKDLYANTVLSGGTTMYPGIADRMQKEITALAPSTMKIKIIAPPERKYSVWIG
TNRC18-ACTB  IMKCDVDIRKDLYANTVLSGGTTMYPGIADRMQKEITALAPSTMKIKIIAPPERKYSVWIG
OAZ1-ACTB    IMKCDVDIRKDLYANTVLSGGTTMYPGIADRMQKEITALAPSTMKIKIIAPPERKYSVWIG
PTMA-ACTB    IMKCDVDIRKDLYANTVLSGGTTMYPGIADRMQKEITALAPSTMKIKIIAPPERKYSVWIG
ELMO1-ACTB   IMKCDVDIRKDLYANTVLSGGTTMYPGIADRMQKEITALAPSTMKIKIIAPPERKYSVWIG
KLF2-ACTB    IMKCDVDIRKDLYANTVLSGGTTMYPGIADRMQKEITALAPSTMKIKIIAPPERKYSVWIG
TPM4-ACTB    IMKCDVDIRKDLYANTVLSGGTTMYPGIADRMQKEITALAPSTMKIKIIAPPERKYSVWIG
LFNG-ACTB    IMKCDVDIRKDLYANTVLSGGTTMYPGIADRMQKEITALAPSTMKIKIIAPPERKYSVWIG

ACTB         GSILASLSTFQQMWISKQEYDESGPSIVHRKCF
B2M-ACTB     GSILASLSTFQQMWISKQEYDESGPSIVHRKCF
FUS-ACTB     GSILASLSTFQQMWISKQEYDESGPSIVHRKCF
MAFK-ACTB    GSILASLSTFQQMWISKQEYDESGPSIVHRKCF
PTBP1-ACTB   GSILASLSTFQQMWISKQEYDESGPSIVHRKCF
ZYX-ACTB     GSILASLSTFQQMWISKQEYDESGPSIVHRKCF
TNRC18-ACTB  GSILASLSTFQQMWISKQEYDESGPSIVHRKCF
OAZ1-ACTB    GSILASLSTFQQMWISKQEYDESGPSIVHRKCF
PTMA-ACTB    GSILASLSTFQQMWISKQEYDESGPSIVHRKCF
ELMO1-ACTB   GSILASLSTFQQMWISKQEYDESGPSIVHRKCF
KLF2-ACTB    GSILASLSTFQQMWISKQEYDESGPSIVHRKCF
TPM4-ACTB    GSILASLSTFQQMWISKQEYDESGPSIVHRKCF
LFNG-ACTB    GSILASLSTFQQMWISKQEYDESGPSIVHRKCF
```

Fig.S2 Alignment of putative protein sequences encoded by 3'-ACTB-fused HFGs a). DDX5-ATP6V0C

```
ATP6V0C       MSESKSGPEYASFFAVMGASAAMVFSALGAAYGTAKSGTGIAAMSVMRPEQIMKSIIPVVM
DDX5-ATP6V0C  -------------------------------------------MSVMRPEQIMKSIIPVVM

ATP6V0C       AGIIAIYGLVVAVLIANSLNDDISLYKSFLQLGAGLSVGLSGLAAGFAIGIVGDAGVRGTA
DDX5-ATP6V0C  AGIIAIYGLVVAVLIANSLNDDISLYKSFLQLGAGLSVGLSGLAAGFAIGIVGDAGVRGTA

ATP6V0C       QQPRLFVGMILILIFAEVLGLYGLIVALILSTK
DDX5-ATP6V0C  QQPRLFVGMILILIFAEVLGLYGLIVALILSTK
```

b). DDX5-CHD2

```
CHD2       MMRNKDKSQEEDSSLHSNASSHSASEEASGSDSGSQSESEQGSDPGSGHGSESNSSSESSE
DDX5-CHD2  MMRNKDKSQEEDSSLHSNASSHSASEEASGSDSGSQSESEQGSDPGSGHGSESNSSSESSE

CHD2       SQSESESESAGSKSQPVLPEAKEKPASKKERIADVKKMWEEYPDVYGVRRSNRSRQEPSRF
DDX5-CHD2  SQSESESESAGSKSQPVLPEAKEKPASKKERIADVKKMWEEYPDVYGVRRSNRSRQEPSRF

CHD2       NIKEEASSGSESGSPKRRGQRQLKKQEKWKQEPSEDEQEQGTSAESEPEQKKVKARRPVPR
DDX5-CHD2  NIKEEASSGSESGSPKRRGQRQLKKQEKWKQEPSEDEQEQGTSAESEPEQKKVKARRPVPR

CHD2       RTVPKPRVKKQPKTQRGKRKKQDSSDEDDDDDEAPKRQTRRRAAKNVSYKEDDDFETDSDD
DDX5-CHD2  RTVPKPRVKKQPKTQRGKRKKQDSSDEDDDDDEAPKRQTRRRAAKNVSYKEDDDFETDSDD

CHD2       LIEMTGEGVDEQQDNSETIEKVLDSRLGKKGATGASTTVYAIEANGDPSGDFDTEKDEGEI
DDX5-CHD2  LIEMTGEGVDEQQDNSETIEKVLDSRLGKKGATGASTTVYAIEANGDPSGDFDTEKDEGEI

CHD2       QYLIKWKGWSYIHSTWESEESLQQQKVKGLKKLENFKKKEDEIKQWYIFHHGLKKCICSLK
DDX5-CHD2  QYLIKWKGWSYIHSTWESEESLQQQKVKGLKKLENFKKKEDEIKQWYIFHHGLKKCICSLK

CHD2         EGKVLK
DDX5-CHD2  EGKVLK
```

c). DDX5-EEF1A1

```
DDX5           MGKEKTHINIVVIGHVDSGKSTTTGHLIYKCGGIDKRTIEKFEKEAAEMGKGSFKYAWVLD
DDX5-EEF1A1  MGKEKTHINIVVIGHVDSGKSTTTGHLIYKCGGIDKRTIEKFEKEAAEMGKGSFKYAWVLD

DDX5         KLKAERERGITIDISLWKFETSKYYVTIIDAPGHRDFIKNMITGTSQADCAVLIVAAGVGE
DDX5-EEF1A1  KLKAERERGITIDISLWKFETSKYYVTIIDAPGHRDFIKNMITGTSQADCAVLIVAAGVGE
```

```
DDX5          FEAGISKNGQTREHALLAYTLGVKQLIVGVNKMDSTEPPYSQKRYEEIVKEVSTYIKKIGY
DDX5-EEF1A1   FEAGISKNGQTREHALLAYTLGVKQLIVGVNKMDSTEPPYSQKRYEEIVKEVSTYIKKIGY

DDX5          NPDTVAFVPISGWNGDNMLEPSANMPWFKGWKVTRKDGNASGTTLLEALDCILPPTRPTDK
DDX5-EEF1A1   NPDTVAFVPISGWNGDNMLEPSANMPWFKGWKVTRKDGNASGTTLLEALDCILPPTRPTDK

DDX5          PLRLPLQDVYKIGGIGTVPVGRVETGVLKPGMVVTFAPVNVTTEVKSVEMHHEALSEALPG
DDX5-EEF1A1   PLRLPLQDVYKIGGIGTVPVGRVETGVLKPGMVVTFAPVNVTTEVKSVEMHHEALSEALPG

DDX5          DNVGFNVKNVSVKDVRRGNVAGDSKNDPPMEAAGFTAQVIILNHPGQISAGYAPVLDCHTA
DDX5-EEF1A1   DNVGFNVKNVSVKDVRRGNVAGDSKNDPPMEAAGFTAQVIILNHPGQISAGYAPVLDCHTA

DDX5          HIACKFAELKEKIDRRSGKKLEDGPKFLKSGDAAIVDMVPGKPMCVESFSDYPPLGRFAVR
DDX5-EEF1A1   HIACKFAELKEKIDRRSGKKLEDGPKFLKSGDAAIVDMVPGKPMCVESFSDYPPLGRFAVR

DDX5          DMRQTVAVGVIKAVDKKAAGAGKVTKSAQKAQKAK
DDX5-EEF1A1   DMRQTVAVGVIKAVDKKAAGAGKVTKSAQKAQKAK
```

d).DDX5-GNAS

```
GNAS        MGVRNCLYGNNMSGQRDIPPEIGEQPEQPPLEAPGAAAPGAGPSPAEEMETEPPHNEPIPV
DDX5-GNAS   ------------------------------------------------------------

GNAS        ENDGEACGPPEVSRPNFQVLNPAFREAGAHGSYSPPPEEAMPFEAEQPSLGGFWPTLEQPG
DDX5-GNAS   ------------------------------------------------------------

GNAS        FPSGVHAGLEAFGPALMEPGAFSGARPGLGGYSPPPEEAMPFEFDQPAQRGCSQLLLQVPD
DDX5-GNAS   ------------------------------------------------------------

GNAS        LAPGGPGAAGVPGAPPEEPQALRPAKAGSRGGYSPPPEETMPFELDGEGFGDDSPPPGLSR
DDX5-GNAS   ------------------------------------------------------------

GNAS        VIAQVDGSSQFAAVAASSAVRLTPAANAPPLWVPGAIGSPSQEAVRPPSNFTGSSPWMEIS
DDX5-GNAS   ------------------------------------------------------------

GNAS        GPPFEIGSAPAGVDDTPVNMDSPPIALDGPPIKVSGAPDKRERAERPPVEEEAAEMEGAAD
DDX5-GNAS   ------------------------------------------------------------

GNAS        AAEGGKVPSPGYGSPAAGAASADTAARAAPAAPADPDSGATPEDPDSGTAPADPDSGAFAA
DDX5-GNAS   ------------------------------------------------------------

GNAS        DPDSGAAPAAPADPDSGAAPDAPADPDSGAAPDAPADPDAGAAPEAPAAPAAAETRAAHVA
DDX5-GNAS   ------------------------------------------------------------

GNAS        PAAPDAGAPTAPAASATRAAQVRRAASAAPASGARRKIHLRPPSPEIQAADPPTPRPTRAS
DDX5-GNAS   ------------------------------------------------------------
```

```
GNAS        AWRGKSESSRGRRVYYDEGVASSDDDSSGDESDDGTSGCLRWFQHRRNRRRRKPQRNLLRN
DDX5-GNAS   -------------------------------------------------------------

GNAS        FLVQAFGGCFGRSESPQPKASRSLKVKKVPLAEKRRQMRKEALEKRAQKRAEKKRSKLIDK
DDX5-GNAS   -------------------------------------------------------------

GNAS        QLQDEKMGYMCTHRLLLLGAGESGKSTIVKQMRILHVNGFNGEGGEEDPQAARSNSDGEKA
DDX5-GNAS   -------------------------------MRILHVNGFNGEGGEEDPQAARSNSDGEKA

GNAS        TKVQDIKNNLKEAIETIVAAMSNLVPPVELANPENQFRVDYILSVMNVPDFDFPPEFYEHA
DDX5-GNAS   TKVQDIKNNLKEAIETIVAAMSNLVPPVELANPENQFRVDYILSVMNVPDFDFPPEFYEHA

GNAS        KALWEDEGVRACYERSNEYQLIDCAQYFLDKIDVIKQADYVPSDQDLLRCRVLTSGIFETK
DDX5-GNAS   KALWEDEGVRACYERSNEYQLIDCAQYFLDKIDVIKQADYVPSDQDLLRCRVLTSGIFETK

GNAS        FQVDKVNFHMFDVGGQRDERRKWIQCFNDVTAIIFVVASSSYNMVIREDNQTNRLQEALNL
DDX5-GNAS   FQVDKVNFHMFDVGGQRDERRKWIQCFNDVTAIIFVVASSSYNMVIREDNQTNRLQEALNL

GNAS        FKSIWNNRWLRTISVILFLNKQDLLAEKVLAGKSKIEDYFPEFARYTTPEDATPEPGEDPR
DDX5-GNAS   FKSIWNNRWLRTISVILFLNKQDLLAEKVLAGKSKIEDYFPEFARYTTPEDATPEPGEDPR

GNAS        VTRAKYFIRDEFLRISTASGDGRHYCYPHFTCAVDTENIRRVFNDCRDIIQRMHLRQYELL
DDX5-GNAS   VTRAKYFIRDEFLRISTASGDGRHYCYPHFTCAVDTENIRRVFNDCRDIIQRMHLRQYELL
```

e). DDX5-HNRNPH3

```
HNRNPH3        MDWVMKHNGPNDASDGTVRLRGLPFGCSKEEIVQFFQGLEIVPNGITLTMDYQGRSTGEAF
DDX5-HNRNPH3   MDWVMKHNGPNDASDGTVRLRGLPFGCSKEEIVQFFQGLEIVPNGITLTMDYQGRSTGEAF

HNRNPH3        VQFASKEIAENALGKHKERIGHRYIEIFRSSRSEIKGFYDPPRRLLGQRPGPYDRPIGGRG
DDX5-HNRNPH3   VQFASKEIAENALGKHKERIGHRYIEIFRSSRSEIKGFYDPPRRLLGQRPGPYDRPIGGRG

HNRNPH3        GYYGAGRGSMYDRMRRGGDGYDGGYGGFDDYGGYNNYGYGNDGFDDRMRDGRGMGGHGYGG
DDX5-HNRNPH3   GYYGAGRGSMYDRMRRGGDGYDGGYGGFDDYGGYNNYGYGNDGFDDRMRDGRGMGGHGYGG

HNRNPH3        AGDASSGFHGGHFVHMRGLPFRATENDIANFFSPLNPIRVHIDIGADGRATGEADVEFVTH
DDX5-HNRNPH3   AGDASSGFHGGHFVHMRGLPFRATENDIANFFSPLNPIRVHIDIGADGRATGEADVEFVTH

HNRNPH3        EDAVAAMSKDKNNMQHRYIELFLNSTPGGGSGMGGSGMGGYGRDGMDNQGGYGSVGRMGMG
DDX5-HNRNPH3   EDAVAAMSKDKNNMQHRYIELFLNSTPGGGSGMGGSGMGGYGRDGMDNQGGYGSVGRMGMG

HNRNPH3        NNYSGGYGTPDGLGGYGRGGGGSGGYYGQGGMSGGGWRGMY
DDX5-HNRNPH3   NNYSGGYGTPDGLGGYGRGGGGSGGYYGQGGMSGGGWRGMY
```

f).DDX5-HNRNPU

```
HNRNPU       MSSSPVNVKKLKVSELKEELKKRRLSDKGLKAELMERLQAALDDEEAGGRPAMEPGNGSLD
DDX5-HNRNPU  ------------------------------------------------------------

HNRNPU       LGGDSAGRSGAGLEQEAAAGGDEEEEEEEEEEEGISALDGDQMELGEENGAAGAADSGPME
DDX5-HNRNPU  ------------------------------------------------------------

HNRNPU       EEEAASEDENGDDQGFQEGEDELGDEEEGAGDENGHGEQQPQPPATQQQQPQQQRGAAKEA
DDX5-HNRNPU  ------------------------------------------------------------

HNRNPU       AGKSSGPTSLFAVTVAPPGARQGQQQAGGKKKAEGGGGGGRPGAPAAGDGKTEQKGGDKKR
DDX5-HNRNPU  ------------------------------------------------------------

HNRNPU       GVKRPREDHGRGYFEYIEENKYSRAKSPQPPVEEEDEHFDDTVVCLDTYNCDLHFKISRDR
DDX5-HNRNPU  ------------------------------------------------------------

HNRNPU       LSASSLTMESFAFLWAGGRASYGVSKGKVCFEMKVTEKIPVRHLYTKDIDIHEVRIGWSLT
DDX5-HNRNPU  -------MESFAFLWAGGRASYGVSKGKVCFEMKVTEKIPVRHLYTKDIDIHEVRIGWSLT

HNRNPU       TSGMLLGEEEFSYGYSLKGIKTCNCETEDYGEKFDENDVITCFANFESDEVELSYAKNGQD
DDX5-HNRNPU  TSGMLLGEEEFSYGYSLKGIKTCNCETEDYGEKFDENDVITCFANFESDEVELSYAKNGQD

HNRNPU       LGVAFKISKEVLAGRPLFPHVLCHNCAVEFNFGQKEKPYFPIPEEYTFIQNVPLEDRVRGP
DDX5-HNRNPU  LGVAFKISKEVLAGRPLFPHVLCHNCAVEFNFGQKEKPYFPIPEEYTFIQNVPLEDRVRGP

HNRNPU       KGPEEKKDCEVVMMIGLPGAGKTTWVTKHAAENPGKYNILGTNTIMDKMMVAGFKKQMADT
DDX5-HNRNPU  KGPEEKKDCEVVMMIGLPGAGKTTWVTKHAAENPGKYNILGTNTIMDKMMVAGFKKQMADT

HNRNPU       GKLNTLLQRAPQCLGKFIEIAARKKRNFILDQTNVSAAAQRRKMCLFAGFQRKAVVVCPKD
DDX5-HNRNPU  GKLNTLLQRAPQCLGKFIEIAARKKRNFILDQTNVSAAAQRRKMCLFAGFQRKAVVVCPKD

HNRNPU       EDYKQRTQKKAEVEGKDLPEHAVLKMKGNFTLPEVAECFDEITYVELQKEEAQKLLEQYKE
DDX5-HNRNPU  EDYKQRTQKKAEVEGKDLPEHAVLKMKGNFTLPEVAECFDEITYVELQKEEAQKLLEQYKE

HNRNPU       ESKKALPPEKKQNTGSKKSNKNKSGKNQFNRGGGHRGRGGFNMRGGNFRGGAPGNRGGYNR
DDX5-HNRNPU  ESKKALPPEKKQNTGSKKSNKNKSGKNQFNRGGGHRGRGGFNMRGGNFRGGAPGNRGGYNR

HNRNPU       RGNMPQRGGGGGGSGGIGYPYPRAPVFPGRGSYSNRGNYNRGGMPNRGNYNQNFRGRGNNR
DDX5-HNRNPU  RGNMPQRGGGGGGSGGIGYPYPRAPVFPGRGSYSNRGNYNRGGMPNRGNYNQNFRGRGNNR

HNRNPU       GYKNQSQGYNQWQQGQFWGQKPWSQHYHQGYY
DDX5-HNRNPU  GYKNQSQGYNQWQQGQFWGQKPWSQHYHQGYY
```

g). DDX5-SF1

```
SF1       MATGANATPLGKLGPPGLPPLPGPKGGFEPGPPPAPGPGAGLLAPGPPPPPPVGSMGALTA
DDX5-SF1  ------------------------------------------------------------
```

```
SF1        AFPFAALPPPPPPPPPPPPQQPPPPPPPPSPGASYPPPQPPPPPPLYQRVSPPQPPPPQPP
DDX5-SF1   -------------------------------------------------------------

SF1        RKDQQPGPAGGGGDFPSKKRKRSRWNQDTMEQKTVIPGMPTVIPPGLTREQERAYIVQLQI
DDX5-SF1   -----------------------------MEQKTVIPGMPTVIPPGLTREQERAYIVQLQI

SF1        EDLTRKLRTGDLGIPPNPEDRSPSPEPIYNSEGKRLNTREFRTRKKLEEERHNLITEMVAL
DDX5-SF1   EDLTRKLRTGDLGIPPNPEDRSPSPEPIYNSEGKRLNTREFRTRKKLEEERHNLITEMVAL

SF1        NPDFKPPADYKPPATRVSDKVMIPQDEYPEINFVGLLIGPRGNTLKNIEKECNAKIMIRGK
DDX5-SF1   NPDFKPPADYKPPATRVSDKVMIPQDEYPEINFVGLLIGPRGNTLKNIEKECNAKIMIRGK

SF1        GSVKEGKVGRKDGQMLPGEDEPLHALVTANTMENVKKAVEQIRNILKQGIETPEDQNDLRK
DDX5-SF1   GSVKEGKVGRKDGQMLPGEDEPLHALVTANTMENVKKAVEQIRNILKQGIETPEDQNDLRK

SF1        MQLRELARLNGTLREDDNRILRPWQSSETRSITNTTVCTKCGGAGHIASDCKFQRPGDPQS
DDX5-SF1   MQLRELARLNGTLREDDNRILRPWQSSETRSITNTTVCTKCGGAGHIASDCKFQRPGDPQS

SF1        AQDKARMDKEYLSLMAELGEAPVPASVGSTSGPATTPLASAPRPAAPANNPPPPSLMSTTQ
DDX5-SF1   AQDKARMDKEYLSLMAELGEAPVPASVGSTSGPATTPLASAPRPAAPANNPPPPSLMSTTQ

SF1        SRPPWMNSGPSESRPYHGMHGGGPGGPGGGPHSFPHPLPSLTGGHGGHPMQHNPNGPPPPW
DDX5-SF1   SRPPWMNSGPSESRPYHGMHGGGPGGPGGGPHSFPHPLPSLTGGHGGHPMQHNPNGPPPPW

SF1        MQPPPPPMNQGPHPPGHHGPPPMDQYLGSTPVGSGVYRLHQGKGMMPPPPMGMMPPPPPPP
DDX5-SF1   MQPPPPPMNQGPHPPGHHGPPPMDQYLGSTPVGSGVYRLHQGKGMMPPPPMGMMPPPPPPP

SF1        SGQPPPPPSGPLPPWQQQQQQPPPPPPPSSSMASSTPLPWQQRSLPAAAMARAMRVRTFRA
DDX5-SF1   SGQPPPPPSGPLPPWQQQQQQPPPPPPPSSSMASSTPLPWQQRSLPAAAMARAMRVRTFRA

SF1        HW
DDX5-SF1   HW
```

h). DDX5-UBB

```
UBB        MQIFVKTLTGKTITLEVEPSDTIENVKAKIQDKEGIPPDQQRLIFAGKQLEDGRTLSDYNI
DDX5-UBB   MQIFVKTLTGKTITLEVEPSDTIENVKAKIQDKEGIPPDQQRLIFAGKQLEDGRTLSDYNI

UBB        QKESTLHLVLRLRGGMQIFVKTLTGKTITLEVEPSDTIENVKAKIQDKEGIPPDQQRLIFA
DDX5-UBB   QKESTLHLVLRLRGGMQIFVKTLTGKTITLEVEPSDTIENVKAKIQDKEGIPPDQQRLIFA

UBB        GKQLEDGRTLSDYNIQKESTLHLVLRLRGGMQIFVKTLTGKTITLEVEPSDTIENVKAKIQ
DDX5-UBB   GKQLEDGRTLSDYNIQKESTLHLVLRLRGGMQIFVKTLTGKTITLEVEPSDTIENVKAKIQ

UBB        DKEGIPPDQQRLIFAGKQLEDGRTLSDYNIQKESTLHLVLRLRGGC
DDX5-UBB   DKEGIPPDQQRLIFAGKQLEDGRTLSDYNIQKESTLHLVLRLRGGC
```

Fig.S3 Analysis of putative proteins encoded by 5'-DDX5-fused HFGs.
a). DDX5-ATP6V0C coded for a truncated ATPase H+ transporting V0 subunit c;
b). DDX5-CHD2 encoded an intact chromodomain helicase DNA binding protein 2;
c).DDX5-EEF1A1 coded for an intact eukaryotic translation elongation factor 1 alpha 1; d).DDX5-GNAS encoded a truncated GNAS complex locus;
e). DDX5-HNRNPH3 coded for an intact heterogeneous nuclear ribonucleoprotein H3; f).DDX5-HNRNPU encoded a truncated heterogeneous nuclear ribonucleoprotein U; g). DDX5-SF1 coded for splicing factor 1;  h). DDX5-UBB encoded an intact ubiquitin B.

```
DDX5              ------------------------------------MSGYSSDRDRGRDRGFGAPRFGGSR
CPSF6-DDX5        ------------------------------------MSAKS-------STRFGAPRFGGSR
TEX2-DDX5         ------------------------------------MRQPA-------DLGFGAPRFGGSR
B2M-DDX5          -------------------------------------------------------------
HNRNPA2B1-DDX5    -------------------------------------------------------------
HNRNPH1-DDX5      -------------------------------------------------------------
PTMA-DDX5         -------------------------------------------------------------
OAZ1-DDX5         MVKSSLQRILNSHCFAREKEGDKPSATIHASRTMPLLSLHS--RGGSSSERFGAPRFGGSR

DDX5              AGPLSGKKFGNPGEKLVKKKWNLDELPKFEKNFYQEHPDLARRTAQEVETYRRSKEITVRG
CPSF6-DDX5        AGPLSGKKFGNPGEKLVKKKWNLDELPKFEKNFYQEHPDLARRTAQEVETYRRSKEITVRG
TEX2-DDX5         AGPLSGKKFGNPGEKLVKKKWNLDELPKFEKNFYQEHPDLARRTAQEVETYRRSKEITVRG
B2M-DDX5          -------------------------------------------------------------
HNRNPA2B1-DDX5    -------------------------------------------------------------
HNRNPH1-DDX5      -------------------------------------------------------------
PTMA-DDX5         -------------------------------------------------------------
OAZ1-DDX5         AGPLSGKKFGNPGEKLVKKKWNLDELPKFEKNFYQEHPDLARRTAQEVETYRRSKEITVRG

DDX5              HNCPKPVLNFYEANFPANVMDVIARQNFTEPTAIQAQGWPVALSGLDMVGVAQTGSGKTLS
CPSF6-DDX5        HNCPKPVLNFYEANFPANVMDVIARQNFTEPTAIQAQGWPVALSGLDMVGVAQTGSGKTLS
TEX2-DDX5         HNCPKPVLNFYEANFPANVMDVIARQNFTEPTAIQAQGWPVALSGLDMVGVAQTGSGKTLS
B2M-DDX5          -------------------MDVIARQNFTEPTAIQAQGWPVALSGLDMVGVAQTGSGKTLS
HNRNPA2B1-DDX5    -------------------MDVIARQNFTEPTAIQAQGWPVALSGLDMVGVAQTGSGKTLS
HNRNPH1-DDX5      -------------------MDVIARQNFTEPTAIQAQGWPVALSGLDMVGVAQTGSGKTLS
PTMA-DDX5         -------------------MDVIARQNFTEPTAIQAQGWPVALSGLDMVGVAQTGSGKTLS
OAZ1-DDX5         HNCPKPVLNFYEANFPANVMDVIARQNFTEPTAIQAQGWPVALSGLDMVGVAQTGSGKTLS


DDX5              YLLPAIVHINHQPFLERGDGPICLVLAPTRELAQQVQQVAAEYCRACRLKSTCIYGGAPKG
CPSF6-DDX5        YLLPAIVHINHQPFLERGDGPICLVLAPTRELAQQVQQVAAEYCRACRLKSTCIYGGAPKG
TEX2-DDX5         YLLPAIVHINHQPFLERGDGPICLVLAPTRELAQQVQQVAAEYCRACRLKSTCIYGGAPKG
B2M-DDX5          YLLPAIVHINHQPFLERGDGPICLVLAPTRELAQQVQQVAAEYCRACRLKSTCIYGGAPKG
HNRNPA2B1-DDX5    YLLPAIVHINHQPFLERGDGPICLVLAPTRELAQQVQQVAAEYCRACRLKSTCIYGGAPKG
HNRNPH1-DDX5      YLLPAIVHINHQPFLERGDGPICLVLAPTRELAQQVQQVAAEYCRACRLKSTCIYGGAPKG
PTMA-DDX5         YLLPAIVHINHQPFLERGDGPICLVLAPTRELAQQVQQVAAEYCRACRLKSTCIYGGAPKG
OAZ1-DDX5         YLLPAIVHINHQPFLERGDGPICLVLAPTRELAQQVQQVAAEYCRACRLKSTCIYGGAPKG


DDX5              PQIRDLERGVEICIATPGRLIDFLECGKTNLRRTTYLVLDEADRMLDMGFEPQIRKIVDQI
CPSF6-DDX5        PQIRDLERGVEICIATPGRLIDFLECGKTNLRRTTYLVLDEADRMLDMGFEPQIRKIVDQI
TEX2-DDX5         PQIRDLERGVEICIATPGRLIDFLECGKTNLRRTTYLVLDEADRMLDMGFEPQIRKIVDQI
B2M-DDX5          PQIRDLERGVEICIATPGRLIDFLECGKTNLRRTTYLVLDEADRMLDMGFEPQIRKIVDQI
HNRNPA2B1-DDX5    PQIRDLERGVEICIATPGRLIDFLECGKTNLRRTTYLVLDEADRMLDMGFEPQIRKIVDQI
HNRNPH1-DDX5      PQIRDLERGVEICIATPGRLIDFLECGKTNLRRTTYLVLDEADRMLDMGFEPQIRKIVDQI
PTMA-DDX5         PQIRDLERGVEICIATPGRLIDFLECGKTNLRRTTYLVLDEADRMLDMGFEPQIRKIVDQI
OAZ1-DDX5         PQIRDLERGVEICIATPGRLIDFLECGKTNLRRTTYLVLDEADRMLDMGFEPQIRKIVDQI
```

```
DDX5             RPDRQTLMWSATWPKEVRQLAEDFLKDYIHINIGALELSANHNILQIVDVCHDVEKDEKLI
CPSF6-DDX5       RPDRQTLMWSATWPKEVRQLAEDFLKDYIHINIGALELSANHNILQIVDVCHDVEKDEKLI
TEX2-DDX5        RPDRQTLMWSATWPKEVRQLAEDFLKDYIHINIGALELSANHNILQIVDVCHDVEKDEKLI
B2M-DDX5         RPDRQTLMWSATWPKEVRQLAEDFLKDYIHINIGALELSANHNILQIVDVCHDVEKDEKLI
HNRNPA2B1-DDX5   RPDRQTLMWSATWPKEVRQLAEDFLKDYIHINIGALELSANHNILQIVDVCHDVEKDEKLI
HNRNPH1-DDX5     RPDRQTLMWSATWPKEVRQLAEDFLKDYIHINIGALELSANHNILQIVDVCHDVEKDEKLI
PTMA-DDX5        RPDRQTLMWSATWPKEVRQLAEDFLKDYIHINIGALELSANHNILQIVDVCHDVEKDEKLI
OAZ1-DDX5        RPDRQTLMWSATWPKEVRQLAEDFLKDYIHINIGALELSANHNILQIVDVCHDVEKDEKLI

DDX5             RLMEEIMSEKENKTIVFVETKRRCDELTRKMRRDGWPAMGIHGDKSQQERDWVLNEFKHGK
CPSF6-DDX5       RLMEEIMSEKENKTIVFVETKRRCDELTRKMRRDGWPAMGIHGDKSQQERDWVLNEFKHGK
TEX2-DDX5        RLMEEIMSEKENKTIVFVETKRRCDELTRKMRRDGWPAMGIHGDKSQQERDWVLNEFKHGK
B2M-DDX5         RLMEEIMSEKENKTIVFVETKRRCDELTRKMRRDGWPAMGIHGDKSQQERDWVLNEFKHGK
HNRNPA2B1-DDX5   RLMEEIMSEKENKTIVFVETKRRCDELTRKMRRDGWPAMGIHGDKSQQERDWVLNEFKHGK
HNRNPH1-DDX5     RLMEEIMSEKENKTIVFVETKRRCDELTRKMRRDGWPAMGIHGDKSQQERDWVLNEFKHGK
PTMA-DDX5        RLMEEIMSEKENKTIVFVETKRRCDELTRKMRRDGWPAMGIHGDKSQQERDWVLNEFKHGK
OAZ1-DDX5        RLMEEIMSEKENKTIVFVETKRRCDELTRKMRRDGWPAMGIHGDKSQQERDWVLNEFKHGK

DDX5             APILIATDVASRGLDVEDVKFVINYDYPNSSEDYIHRIGRTARSTKTGTAYTFFTPNNIKQ
CPSF6-DDX5       APILIATDVASRGLDVEDVKFVINYDYPNSSEDYIHRIGRTARSTKTGTAYTFFTPNNIKQ
TEX2-DDX5        APILIATDVASRGLDVEDVKFVINYDYPNSSEDYIHRIGRTARSTKTGTAYTFFTPNNIKQ
B2M-DDX5         APILIATDVASRGLDVEDVKFVINYDYPNSSEDYIHRIGRTARSTKTGTAYTFFTPNNIKQ
HNRNPA2B1-DDX5   APILIATDVASRGLDVEDVKFVINYDYPNSSEDYIHRIGRTARSTKTGTAYTFFTPNNIKQ
HNRNPH1-DDX5     APILIATDVASRGLDVEDVKFVINYDYPNSSEDYIHRIGRTARSTKTGTAYTFFTPNNIKQ
PTMA-DDX5        APILIATDVASRGLDVEDVKFVINYDYPNSSEDYIHRIGRTARSTKTGTAYTFFTPNNIKQ
OAZ1-DDX5        APILIATDVASRGLDVEDVKFVINYDYPNSSEDYIHRIGRTARSTKTGTAYTFFTPNNIKQ

DDX5             VSDLISVLREANQAINPKLLQLVEDRGSGRSRGRGGMKDDRRDRYSAGKRGGFNTFRDREN
CPSF6-DDX5       VSDLISVLREANQAINPKLLQLVEDRGSGRSRGRGGMKDDRRDRYSAGKRGGFNTFRDREN
TEX2-DDX5        VSDLISVLREANQAINPKLLQLVEDRGSGRSRGRGGMKDDRRDRYSAGKRGGFNTFRDREN
B2M-DDX5         VSDLISVLREANQAINPKLLQLVEDRGSGRSRGRGGMKDDRRDRYSAGKRGGFNTFRDREN
HNRNPA2B1-DDX5   VSDLISVLREANQAINPKLLQLVEDRGSGRSRGRGGMKDDRRDRYSAGKRGGFNTFRDREN
HNRNPH1-DDX5     VSDLISVLREANQAINPKLLQLVEDRGSGRSRGRGGMKDDRRDRYSAGKRGGFNTFRDREN
PTMA-DDX5        VSDLISVLREANQAINPKLLQLVEDRGSGRSRGRGGMKDDRRDRYSAGKRGGFNTFRDREN
OAZ1-DDX5        VSDLISVLREANQAINPKLLQLVEDRGSGRSRGRGGMKDDRRDRYSAGKRGGFNTFRDREN

DDX5             YDRGYSSLLKRDFGAKTQNGVYSAANYTNGSFGSNFVSAGIQTSFRTGNPTGTYQNGYDST
CPSF6-DDX5       YDRGYSSLLKRDFGAKTQNGVYSAANYTNGSFGSNFVSAGIQTSFRTGNPTGTYQNGYDST
TEX2-DDX5        YDRGYSSLLKRDFGAKTQNGVYSAANYTNGSFGSNFVSAGIQTSFRTGNPTGTYQNGYDST
B2M-DDX5         YDRGYSSLLKRDFGAKTQNGVYSAANYTNGSFGSNFVSAGIQTSFRTGNPTGTYQNGYDST
HNRNPA2B1-DDX5   YDRGYSSLLKRDFGAKTQNGVYSAANYTNGSFGSNFVSAGIQTSFRTGNPTGTYQNGYDST
```

| | |
|---|---|
| HNRNPH1-DDX5 | YDRGYSSLLKRDFGAKTQNGVYSAANYTNGSFGSNFVSAGIQTSFRTGNPTGTYQNGYDST |
| PTMA-DDX5 | YDRGYSSLLKRDFGAKTQNGVYSAANYTNGSFGSNFVSAGIQTSFRTGNPTGTYQNGYDST |
| OAZ1-DDX5 | YDRGYSSLLKRDFGAKTQNGVYSAANYTNGSFGSNFVSAGIQTSFRTGNPTGTYQNGYDST |

| | |
|---|---|
| DDX5 | QQYGSNVPNMHNGMNQQAYAYPATAAAPMIGYPMPTGYSQ |
| CPSF6-DDX5 | QQYGSNVPNMHNGMNQQAYAYPATAAAPMIGYPMPTGYSQ |
| TEX2-DDX5 | QQYGSNVPNMHNGMNQQAYAYPATAAAPMIGYPMPTGYSQ |
| B2M-DDX5 | QQYGSNVPNMHNGMNQQAYAYPATAAAPMIGYPMPTGYSQ |
| HNRNPA2B1-DDX5 | QQYGSNVPNMHNGMNQQAYAYPATAAAPMIGYPMPTGYSQ |
| HNRNPH1-DDX5 | QQYGSNVPNMHNGMNQQAYAYPATAAAPMIGYPMPTGYSQ |
| PTMA-DDX5 | QQYGSNVPNMHNGMNQQAYAYPATAAAPMIGYPMPTGYSQ |
| OAZ1-DDX5 | QQYGSNVPNMHNGMNQQAYAYPATAAAPMIGYPMPTGYSQ |

Fig.S4 Alignments of putative protein sequences encoded by 3'-DDX5-fused HFGs.

```
KLF2           ------------------------------M---ALSEPI--LPSFSTFASPC-------
B2M-KLF2       --------------------------MSRSV---ALAVLA--LLSLS-------------
EPS15L1-KLF2   --------------------------MRGPVYGPALSRDRRPAHRPPVVPVRTRGCSPAP
TPM4-KLF2      --------------------------MAGLN---SLEAVKRKIQALQQQADEAEDRAQGL
ACTB-KLF2      ---------------------------------LWLERPRRRPIKPSGATRHHRRDRVRP
AKAP8-KLF2     ------------------------------------------------------------
ELL-KLF2       MAALKEDRSYGLSCGRVSDGSKVSVFHVKLTDSALRAFESYRARQALPLQLGRLRLEVCAL
PTBP1-KLF2     ---------------------------------MGSARGGRVRASPPFCESITRSRWVGS

KLF2           -------RERGL---------QEVRSPTTATGTAAAGSLRAQTSSRATTESTRATGHSSAI
B2M-KLF2       ----------GL---------EAIQRPTTATGTAAAGSLRAQTSSRATTESTRATGHSSAI
EPS15L1-KLF2   VPPALGARVRGKMAAPLIPLSQQVRSPTTATGTAAAGSLRAQTSSRATTESTRATGHSSAI
TPM4-KLF2      QRELDGERERR----------EKVRSPTTATGTAAAGSLRAQTSSRATTESTRATGHSSAI
ACTB-KLF2      A--STEPRLCRSAARPH-PPPGEKPYHCNWDGCG----WKFARSDELTRHYRKHTGHRPFQ
AKAP8-KLF2     --------------MDQ-GYGGEKPYHCNWDGCG----WKFARSDELTRHYRKHTGHRPFQ
ELL-KLF2       RR--AHAPLPKAHGPPAIPV-----------PSVRSCLLALRSPGAAHETAHVAGTPPPT
PTBP1-KLF2     C--YSGASTPSPAGLLCVPWTGEKPYHCNWDGCG----WKFARSDELTRHYRKHTGHRPFQ

KLF2           CAIVPSRAPITWRCT-------------------------
B2M-KLF2       CAIVPSRAPITWRCT-------------------------
EPS15L1-KLF2   CAIVPSRAPITWRCT-------------------------
TPM4-KLF2      CAIVPSRAPITWRCT-------------------------
ACTB-KLF2      CHLCDRAFSRSDHLALHMKRHM------------------
AKAP8-KLF2     CHLCDRAFSRSDHLALHMKRHM------------------
ELL-KLF2       CAR-PWRVPRAGRGPLPNCDWYLLDPENRAGHSVATEGLPR
PTBP1-KLF2     CHLCDRAFSRSDHLALHMKRHM------------------
```

Fig.S5 Alignment of putative protein sequences encoded by 3'-KLF2-fused HFGs.
